# Gene Expression Distribution Deconvolution in Single Cell RNA Sequencing

**DOI:** 10.1101/227033

**Authors:** Jingshu Wang, Mo Huang, Eduardo Torre, Hannah Dueck, Sydney Shaffer, John Murray, Arjun Raj, Mingyao Li, Nancy R. Zhang

## Abstract

Single-cell RNA sequencing (scRNA-seq) enables the quantification of each gene’s expression distribution across cells, thus allowing the assessment of the dispersion, burstiness, and other aspects of its distribution beyond the mean. These statistical characterizations of the gene expression distribution are critical for understanding expression variation and for selecting marker genes for population heterogeneity. However, scRNA-seq data is noisy, with each cell typically sequenced at low coverage, thus making it difficult to infer properties of the gene expression distribution from raw counts. Based on a re-examination of 9 public data sets, we propose a simple technical noise model for scRNA-seq data with Unique Molecular Identifiers (UMI). We develop DESCEND, a method that deconvolves the true cross-cell gene expression distribution from observed scRNA-seq counts, leading to improved estimates of properties of the distribution such as dispersion and burstiness. DESCEND can adjust for cell-level covariates such as cell size, cell cycle and batch effects. DESCEND’s noise model and estimation accuracy are further evaluated through comparisons to RNA FISH data, through data splitting and simulations, and through its effectiveness in removing known batch effects. We demonstrate how DESCEND can clarify and improve downstream analyses such as finding differentially bursty genes, identifying cell types, and selecting differentiation markers.

## Introduction

Cells are the basic biological units of multicellular organisms. Within a cell population, individual cells vary in their gene expression levels, reflecting the dynamics of transcription across cells [51, 42, 55, 33, 50]. Traditional microarray and bulk RNA-seq technologies profile the average gene expression level of all cells in the population. In contrast, recent single cell RNA-seq (scRNA-seq) methods enable the quantification of a much richer set of features of the gene expression distribution across cells. For example, measures of dispersion such as coefficient of variation (CV) and Gini coefficients can be used to elucidate biological states that are not reflected in the population average [49, 28, 62, 48]. Measures of expression burstiness, alternatively, allow a better understanding of transcriptional regulation at the single cell level [22, 50].

However, it is challenging to compute such distribution based statistics of true gene expression due to the technical noise in scRNA-seq data [9, 56, 25, 30, 53]. Single cell RNA sequencing protocols are complex, involving multiple steps each contributing to the substantially increased noise level of scRNA-seq relative to bulk RNA-seq. Unique Molecular Identifiers (UMI) [27] were introduced as a barcoding technique to reduce amplification noise, but due to the low efficiency that plagues most single cell experiments, the observed expression distribution computed from observed UMI counts is, for most genes, still a poor representation of its true expression distribution. Even the simple sampling variability in the experiment can distort distribution measurement such as dispersion and zero-inflation of the observed counts largely from that of the true expression.

Recently, many computational methods for scRNA-seq analysis have been proposed, including methods for quantifying dispersion, characterization of transcriptional bursting, and finding differentially expressed genes [24, 12, 16, 3, 23, 41, 28, 38, 31]. Though some of these work has taken the technical noise into consideration, to our knowledge, there is currently no method for recovering the entire cross-cell gene expression distribution from scRNA-seq data, nor for comparing distribution features beyond the mean while controlling for cell-level factors such us cell cycle and cell size. In addition, there is still a lack of thorough analysis of technical noise model that can properly fit scRNA-seq data.

Here we develop DESCEND (DEconvolution of Single Cell ExpressioN Distribution), a statistical method that deconvolves the true cross-cell gene expression distribution from observed scRNA-seq counts and quantifies the dependence between features of this distribution and cell-level covariates such as cell size and cell type. DESCEND adopts the “G-modeling” empirical Bayes distribution deconvolution framework [10], which avoids constraining parametric assumptions. The accuracy of DESCEND is evaluated using RNA FISH data generated from the same cell population [56], and is further assessed through sample splitting and parametric simulations. Our evaluations show that, under very reasonable data quality assumptions, DESCEND can accurately deconvolve the true gene expression distribution, leading to improved characterization of dispersion and expression burstiness. We show through case studies how these improved estimates lead to more accurate downstream analyses such as cell type classification, marker gene selection, and differential expression analysis. We benchmark against existing methods [34, 28, 38] on specific analyses in which comparable methods exist, but focus on novel applications of DESCEND in the case studies.

Although the DESCEND framework can be used with any technical noise, the data sets we use in this paper all employ UMI, for which we have a clear understanding of the noise model. There has been a lot of debate regarding what noise model to use, even for UMI-based scRNA-seq data. However, through a re-analysis of nine public data sets with UMI, we show that a Poisson distribution is sufficient to capture the technical noise in single cell UMI counts, once the underlying biological variations and cross-cell differences in library size have been accounted for. Given this result, DESCEND adopts the Poisson noise model for single cell UMI counts as its default setting, thus achieving fast computation and stable estimation. We show using the data from Tung et al. [57] that, with this noise model, DESCEND can effectively remove artificial differences between known experimental batches.

We demonstrate the applications of DESCEND in three case studies. The first is an analysis of gene expression burstiness. Ample evidence from RNA FISH studies have shown that, for most genes, true single cell expression is not Poisson, even in a seemingly homogeneous population. This observed cell-level heterogeneity in RNA count is, in part, due to the bursty process of gene transcription, where periods of RNA synthesis is followed by periods of inactivity [5, 43, 6]. The Beta-Poisson distribution has been proposed to account for the inflation of zeros and long tail of this bursty RNA count distribution in RNA FISH data. For scRNA-seq data, accounting for technical noise is especially critical in fitting such mixture models, as most genes in any given cell have zero or low observed count, due both to biological inactivity and to technical noise [50, 1, 8]. RNA FISH experiments have also been used to explore the dependence of bursting on cell size, cell cycle, and mean expression level [39]. Yet these experiments were performed on a limited set of genes in an in vitro setting. There has not been a transcriptome-wide exploration of such relationships by scRNA-seq in real tissues, partly due to the lack of an effective statistical and computational framework.

We use DESCEND to characterize the relationship between expression burstiness and cell size and to detect differential burstiness between cell types using the mouse brain scRNA-seq data from Zeisel et al. [62]. Burstiness is quantified by two parameters: nonzero fraction (fraction of cells where the gene is expressed) and nonzero mean (mean expression level among cells with positive expression). We show that, transcriptome-wide, in all cell types analyzed, cell size is positively correlated with nonzero fraction and may have a sub-linear relationship with nonzero mean. These findings are replicated in multiple cell types and in RNA-FISH data of a human melanoma cell line [56], suggesting that the relationship is real and may be widespread. Hence, in detecting differences in expression burstiness across cell types or across conditions, one needs to account for cell size to avoid confounding. We illustrate how such an analysis can be done using DESCEND.

Our second and third case studies are on the dispersion of gene expression. Precise estimates of expression dispersion, through quantities such as CV, Gini coefficient and Fano factors, form the basis of many single cell analysis pipelines. For example, a fundamental recurring analysis in scRNA-seq is the ranking of genes by dispersion to select informative genes for cell type clustering. Beyond its utility in gene pre-filtering, transcriptional variability has also long been recognized to be intrinsically important to fundamental biological processes [11, 32, 14]. For example, a recent study shows that cell-to-cell transcriptional variability is related to aging [36]. Dispersion measures such as Gini coefficient and CV computed on raw scRNA-seq data are severely biased due to zero inflation, as shown by Torre et al. [56] and Klein et al. [28]. Although Klein et al. [28] derived a formula to correct the CV computed from raw counts for technical variation, no comparable method exists for the Gini coefficient, the latter, as we show, is a more robust measure of dispersion. Comparisons to FISH measurements taken from the same cell population establish that DESCEND provides unbiased and robust estimates of CV and Gini coefficients for the true gene expression distribution. We demonstrate how such improved dispersion estimates lead to better marker gene selection and cell type identification.

In summary, we developed DESCNED to deconvolve the true cross-cell gene expression distri-bution from UMI-based scRNA-seq data. One key contribution of DESCEND is the validation of a simple noise model of the UMI counts using public data sets, which leads to good performance of DESCEND in batch effect removal and when compared with RNA FISH. After confirming the accuracy of DESCEND, we then illustrate the utility of the deconvolved distribution in three case studies. We demonstrate that this framework allows better characterization of expression burstiness and population heteogeneity, which is necessary for quantifying the stochasticity of gene expression across cells.

## Results

### Model Overview

Figure 1 gives an overview of the DESCEND framework. The observed counts in an scRNA-seq experiment is a noisy reflection of true expression levels. We model the observed count *Y_cg_* for gene *g* in cell *c* as a convolution of the true gene expression *λ_cg_* and technical noise,

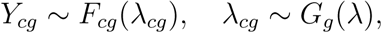

where *F_cg_*(·) quantifies technical noise and *G_g_* represents the true expression distribution of gene *g* across cells. DESCEND deconvolves **G_g_** from the noisy observed counts *Y_cg_*, thus allows for estimation of any distribution-related quantity of interest. One difference between DESCEND and pre-existing methods is that DESCEND models *G_g_* using a spline-based exponential family, which avoids restrictive parametric assumptions while allowing the flexible modeling of covariate effects.

**Figure 1:**
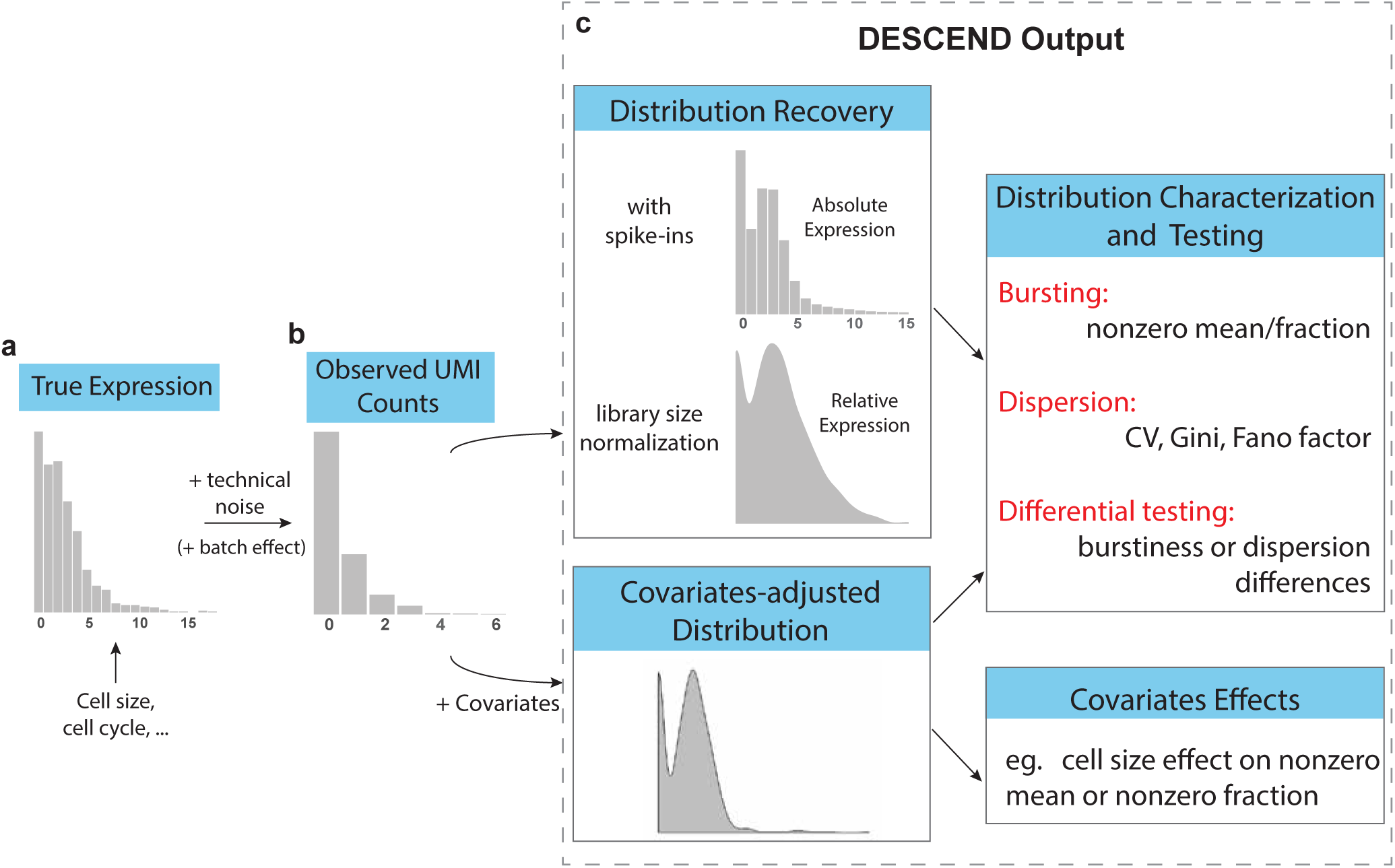
Illustration of the framework. (**b**)The cross-cell distribution of observed counts *Y_cg_* is assumed to be a convolution of (**a**)the distribution of true gene expression and technical noise. (**c**)For each gene, the output of DESCEND includes: the distribution of the absolute expression levels when cell-specific efficiency constants can be estimated from ERCC spike-ins, the distribution of relative expression with library size normalization when spike-in genes are not available, the distribution of covariates-adjusted expression level if covariates are presented, estimates of the burstiness and dispersion parameters, differential testing results comparing the change between two cell populations, and the effects of covariates on gene expression when recovering the covariates-adjusted distribution.

Currently, DESCEND focuses on single cell experiments that utilize UMI. For extension to non-UMI read counts, see Discussion. In the next section, we show through a re-examination of public data that the Poisson distribution is sufficient for capturing the technical noise in UMI counts, after accounting for cross-cell differences in library size. Thus, for UMI-based single cell RNA-seq data, DESCEND employs the noise model *Y_cg_* ∼ Poisson(*α_c_λ_cg_*), where *α_c_* is a cell specific scaling constant. By default, DESCEND sets *α_c_* to be the total UMI count of cell *c*, which leads to the interpretation of *λ_cg_* being the relative expression of gene *g* in the cell. If reliable cell-specific spike-ins are available, one could compute the efficiency, defined as the proportion of transcripts in the cell that are sequenced, and set *α_c_* to the efficiency of cell *c*. This latter definition leads to the interpretation of *λ_cg_* being the absolute expression of gene *g* in the cell.

The true gene expression distributions *G_g_* are expected to be complex, owing to the possibility of multiple cell sub-populations and to the transcriptional heterogeneity within each sub-population. In particular, this distribution may have several modes and an excessive amount of zeros, and can not be assumed to abide by known parametric forms. To allow for such complexity, DESCEND adopts the G-modeling empirical Bayes distribution deconvolution technique in Efron [10], and models the gene expression distribution as a zero-inflated exponential family distribution, which has the zero-inflated Poisson, logNormal and Gamma distributions as special cases. The G-modeling technique uses natural cubic spline functions to estimate the shape of the gene expression distribution adaptively from the observed counts (see Methods). Model complexity and estimation accuracy are balanced by discretization of the gene expression distribution and adding shrinkage penalties to the likelihood, as suggested in Efron [10].

One meaningful characteristic of the gene expression distribution *G_g_* is the proportion of cells where the gene has non-zero expression, that is,

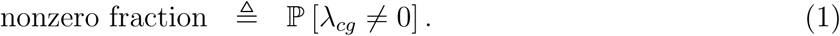

Complementary to the nonzero fraction is the nonzero mean, defined as the average expression level among cells where the gene has nonzero expression,

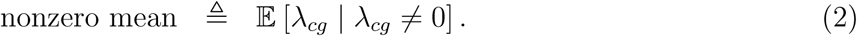

The concepts of nonzero fraction and nonzero mean have appeared, under varying definitions and differing names, in single cell studies [37, 50, 38], yet many existing approaches to estimate them [37, 24, 50, 7, 60] do not account for technical noise. If the population from which the cells are sampled can be assumed to be ergodic, then a two-state transcriptional bursting model [22, 43, 21], formulated as a periodic stochastic dynamic process, leads to a Beta-Poisson distribution for gene expression across cells. In that scenario, Equations (1) and (2) will be highly correlated with the burst frequency and burst size parameters defined in the Beta-Poisson distribution. However, the strong ergodicity assumption is, in most cases, too ideal for scRNA-seq experiments, in which cell populations are unavoidably heterogeneous even when limited to a specific cell type. In DESCEND, we choose not to assume the Beta-Poisson distribution, which reduces estimation complexity and allows more flexibility.

DESCEND allows the inclusion of covariate effects on both the nonzero fraction and nonzero mean. When covariates are specified, DESCEND uses a log linear model for the covariates effect on nonzero mean and a logit model for the covariate effect on nonzero fraction. Thus, when covariates are specified, the deconvolution result is the covariate-adjusted distribution of gene expression, see Methods for details.

DESCEND also computes standard errors and performs hypotheses tests on distribution features such as dispersion and expression burstiness parameters. See Methods for details.

## Model Assessment and Validation

### Technical noise model for UMI-based scRNA-seq experiments

For UMI-based scRNA-seq data, Kim et al. [25] gave an analytic argument for a Poisson error model, which we discussed and clarify further in Methods. Several studies [2, 24, 15] used spike in sets and bulk RNA splitting experiments to explore the technical noise in scRNA-seq data, finding that a Poisson distribution for UMI-based counts is plausible, but raised the issue of over-dispersion. While the analyses from these studies were insightful, we believe that they failed to account for the inevitable random variations across cells/samples in the input of spike-in when splitted at low concentrations. We re-examined the spike-in data from nine UMI-based scRNA-seq datasets, including seven different scRNA-seq protocols (Figure 2a). The UMI data sets of ERCC we analyzed are from Jaitin et al. [19], Zeisel et al. [62], Klein et al. [28], Macosko et al. [35], Hashimshony et al. [18], Tung et al. [57], Zheng et al. [63] and Svensson et al. [54]. All the data sets except for Tung et al. [57] have also been analyzed in Svensson et al. [54], who showed that capture efficiency vary substantially across cells within each experiment, and between experimental protocols. We show that after accounting for the cell-to-cell variation in efficiency, the technical noise of UMI counts is simply Poisson for most datasets.

**Figure 2:**
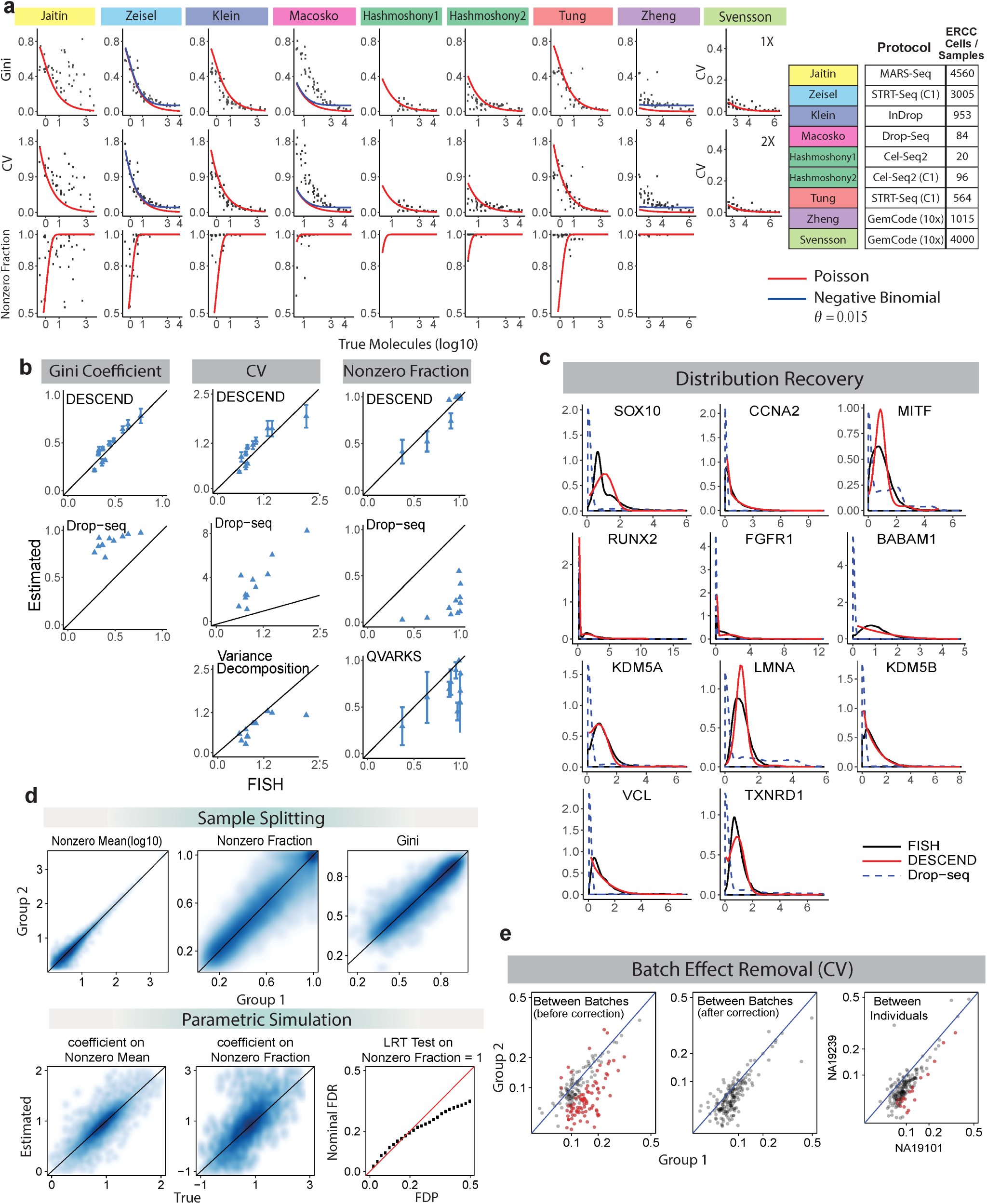
Validation of DESCEND. (**a**)Noise model: the Poisson technical noise model is tested using 9 different ERCC datasets that has UMI. Svensson et al. [54] has two ERCC datasets with different concentrations (1X and 2X, 2000 cells each). Each dot is for a spike-in gene. (**b**)RNA FISH: the Gini coefficients, CV and the Nonzero Fraction for 11 profiled genes are compared between the smRNA FISH values and the DESCEND estimated values from Drop-seq data [56]. Values computed directly from Drop-seq counts and using other methods are also included for comparison. (**c**)FISH distribution recovery: the deconvolved true gene relative expression distribution is compared among smRNA FISH distribution, DESCEND and the distribution of Drop-seq normalized counts. (**d**)For the sample splitting experiments, distribution based measurements are compared between the two split groups. For the parametric simulation, coefficients of the cell size covariates are compared against the true values and the nominal FDR and actual false discovery proportion (FDP) for the likelihood ratio test is compared. (**e**)Batch effect removal [57]: the DESCEND estimated CV are compared between two artificial groups before (left) and after (middle) adding batches as covariates. Each dot is a gene. The estimated CV after adding batches as covariates are also compared between two individuals(right). The red dots are significantly differentially expressed genes (of CV) when FDR in controlled at level 5%.

For ERCC spike-in “genes”, the observed count for each gene in each cell depends on the number of input molecules and the technical noise. Due to the low spike-in concentration added to each cell, the number of input molecules for each spike-in is not fixed, but random with a certain target expectation. If we assume that the molecules in the spike-in dilution are randomly dispersed then the number that result in each cell partition is Poisson with mean computable from the dilution ratios (see Supplementary Text for more details). If the molecules in the spike-in dilution are not randomly dispersed, e.g. due to clumping, or if there are uncontrolled batch issues, then the input number of spike-in molecules for each cell would be over-dispersed compared to the Poisson.

The key observation here is that the input quantity of spike-in molecules is not fixed across cells, as assumed by current studies, but random with Poisson noise in the ideal case of perfect random dispersion with no batch variation. Such randomness of input molecule should not be counted as the technical noise of scRNA-seq experiments, as they are not present in the real biological genes. Previous analyses of spike-in data have attributed this level of spike-in variation to the technical noise of the experiment, thus inflating their estimates of technical noise dispersion.

To assess whether the technical noise of each scRNA-seq data set is Poisson after accounting for cell-specific efficiency, we performed the following analysis: DESCEND is applied to each spike-in gene in each data set with the error model

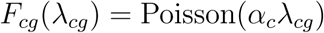

to obtain the underlying distribution of the input molecule counts. If this model is a good approximation to the true technical noise distribution of the scRNA-seq experiment, *and* if the spike-ins are ideal in the sense described above, then the DESCEND recovered input molecule (*λ_cg_*) distributions of the spike-in genes should be Poisson. Conversely, if the recovered distributions show zero inflation or over-dispersion as compared to the Poisson distribution, then that may be due to either a mis-specified technical noise model or to unaccounted experimental factors in the spike-ins. Note that the application of DESCEND does not require spike-ins, here, the spike-ins from these nine studies are simply used to assess whether the technical noise model assumed by DESCEND is appropriate.

Figure 2a shows that the DESCEND recovered distribution in all but one (Macosko et al. [35]) of the nine UMI datasets have over-dispersion *θ* < 0.015 as compared to the Poisson, where *θ* is defined in the variance-mean equation σ^2^ = *μ* + *θμ*^2^. Svensson et al. [54] contains two datasets at different concentration levels and because of the low efficiency (less than 0.01% on average per cell) of the experiment, their dispersion measured by CV is calculated using a simple moment method (more details see Supplementary Text). The over-dispersion is effectively zero in six of the datasets and less than 0.015 in the other two, indicating that the technical noise model used by DESCEND well approximates the technical noise in the data. As discussed above, the mis-fit of the Poisson to the recovered distribution for Macosko et al. [35] data can be either due to a wrong technical noise model or to clumping in the spike-ins. Note that for Macosko et al. [35], the over-dispersion is high for low input values, which is reverse that of typical RNA-seq experiments. This pattern of over-dispersion can be explained by a clumping model on the input molecules. (see Methods for discussion)

### Evaluation of deconvolution accuracy using RNA FISH as gold standard

Next, we evaluate the accuracy of DESCEND on the data from Torre et al. [56], where Drop-seq and RNA FISH are both applied to the same melanoma cell line. 5763 cells and 12241 genes were kept for analysis from the Drop-seq experiment, with median 1473 UMIs per cell. Of these genes, 24 were profiled using RNA FISH (VCF and FOSL1 removed from the original data, see Supplementary Text). We further exclude genes that have zero Drop-seq observed counts in more than 98% of the cells, resulting in 12 genes. The relative gene expression distributions are recovered by DESCEND and are compared with the gene expression profiles using RNA FISH. Since the distribution recovered by DESCEND reflects relative expression levels (a.k.a. concentrations), for comparability the expression of each gene in FISH was normalized by GAPDH [39].

Both CV and Gini coefficients can be accurately recovered using DESCEND (Figure 2b). In comparison, Gini coefficients and CV computed on the original Drop-seq counts, normalized by library size, show very poor correlation and substantial positive bias; this agrees with previous results [28, 56]. For CV, the variance decomposition approach modified from Klein et al. [28] (see Supplementary Text) shows slight bias towards 0 compared with the true CV values calculated from RNA FISH. This analysis also shows that the one standard deviation error bars of DESCEND the fluctuation of our estimates.

DESCEND provides reasonably accurate estimates of the nonzero fraction, despite the low sequencing depth of this data set (Figure 2b). In contrast, the naive estimate, derived from the proportion of nonzero raw counts for each gene, is grossly inflated due to the low sequencing depth and is not a reliable estimator of nonzero fraction. DESCEND outperforms QVARKS [38], a recent method estimating the nonzero fraction (called “ON fraction” in the original paper) using a Bayesian approach.

Finally, DESCEND recovers the shape of the relative gene expression distribution (Figure 2c), as shown by comparison to the FISH data. In comparison, the distribution of the normalized observed counts are quite different from their FISH counterparts, showing severe zero inflation and increased skewness.

### Assessment of estimation accuracy and test validity by sample splitting and parametric simulations

We further evaluate the accuracy of DESCEND on sample-splitting and parametric simulations. For both experiments, we start with the observed counts of one cell type, the oligodendrocyte cells, from Zeisel et al. [62]. The data also contains ERCC spike-ins for every cell, from which we estimate the cell-specific efficiencies.

First, in the sample-splitting experiment, the 820 cells belonging to the cell type are randomly split into two equal-sized groups and, within each group, DESCEND is applied to recover the distribution of the absolute gene expression using the cell-specific efficiencies as *α_c_*. The burstiness and dispersion parameters obtained from the two groups are compared. Since the two groups are obtained by randomly splitting a roughly homogeneous cell population, there should not be any real differences between them. The magnitude of difference between DESCEND estimates across the two groups reflect estimation inaccuracy. We found good agreement in estimates of nonzero mean, nonzero fraction and Gini coefficients between the two groups (Figure 2d). Nonzero fraction is intrinsically more difficult to estimate, and thus vary more substantially between the two groups, but the error is also controlled for most genes.

The sample splitting experiment gives a model free assessment of DESCEND estimation variance, but can not be used to assess estimation bias. To assess estimation bias, we perform a parametric simulation experiment where UMI counts are simulated using parameters estimated from the real data. For details of the simulation, see Methods.

The results of DESCEND on this simulated data set indicate that, under the correct model assumptions, it gives unbiased estimates for the effects of cell size on nonzero fraction and nonzero mean (Figure 2d). Nonzero fraction, CV and the Gini coefficients also get accurate and unbiased estimates (Figure S1a). The nonzero fraction is harder to estimate in a covariates-adjusted model, owing to the fact that the covariates-adjusted distribution is no longer a count-based distribution, but continuous, making it more difficult to separate zero from non-zero but small values. Despite this barrier, the nonzero fraction estimates for most genes are reliable. Finally, with Benjamini-Hochberg procedure, DESCEND effectively controls the FDR in the test of whether the nonzero fraction is 1 (Figure 2d).

### Batch effects can be removed in differential analysis by adding batch as covariate

Tung et al. [57] performed scRNA-seq on three human iPSC cell lines, three technical replicates per cell line, and showed that there can be substantial variation between technical replicates. They referred to the variation between technical replicates as the “batch effect”. Tung et al. [57] further showed that simple ERCC spike-in adjustment and library size normalization can not effectively remove the technical “batch effect”, and proposed a regression-based method. We apply DESCEND to this data to see if using batch as a covariate in DESCEND effectively removes technical differences between replicates.

Starting from the data of Tung et al. [57], we created two groups of cells, each containing 150 cells obtained by pooling 50 cells randomly selected from each of the three individuals. Thus, the two groups of cells should have no biological differences. However, the replicates (batches) are manually chosen to manifest the technical batch effect between the two groups: The first group contains cells sampled from one replicate for each subject: NA19098 replicate 1, NA19101 replicate 2, and NA19239 replicate 1; the second group contains cells sampled from another replicate from each subject: NA19098 replicate 3, NA19101 replicate 1, and NA19239 replicate 2. With the two groups of cells constructed in this way, any detection made during differential testing must be a false positive due to the technical differences between replicates (batch effects).

DESCEND was applied to this data to test for differences in CV and Gini coefficient between the two groups (Figure 2e, Figure S1b). Without consideration of batch, DESCEND indeed finds many (false positive) differences in CV and Gini coefficients. However, with batches added as covariates in the DESCEND model, the dispersion estimates from the two groups are comparable, and no significant detections are made. The fact that spike-in based normalization can not effectively remove this batch effect, which is effectively removed by DESCEND model, indicates that the technical differences between batches are gene specific. We also conducted differential dispersion analysis using DESCEND between two biologically different samples (the three replicates from NA19101 versus the three replicates from NA19239), with batch as covariate, and found significant changes in dispersion. The fact that significant differences are found between biologically different samples, but not between biologically identical samples, suggests that DESCEND effectively removes the batch effect while preserving biological signal.

## Case Studies

### Differential testing of nonzero fraction and mean, accounting for differences in cell size

At the single cell level, most genes are bursty, being inactive in some cells and active in other cells. In Equations (1)-(2), we defined the nonzero fraction and nonzero mean to characterize the burstiness of the gene expression distribution across cells. scRNA-seq allow the detection of changes in these burstiness parameters across cell populations. However, such analyses are easily confounded by not only technical noise, but also cell-level covariates such as cell size. Using DESCEND, we analyze the scRNA-seq data of mouse hippocampal region from Zeisel et al. [62], where the 3005 cells are classified into 7 major cell types. Our goal is to characterize the dependence of nonzero fraction and nonzero mean on cell size, and to compare expression burstiness patterns across cell types, controlling for cell size.

First, consider the transcriptome-wide patterns of expression burstiness, without adjusting for cell size. We applied DESCEND to each gene in each cell type separately, with no added covariates, see Supplementary Text. As shown in Figure 3a, the deconvolved gene expression distributions for most genes have much larger nonzero fraction in the neuron cell types (CA1 pyramidal, S1 pyramidal and Interneurons) as compared to the non-neuron cell types (astrocytes-ependymal, endothelial-mural, microglia and oligodentrocytes), thus suggesting that gene expression in neurons are, in general, less bursty. However, neurons are known to be larger cells, and in this data, the cell size estimates are substantially larger for the neurons as compared to the non-neurons (Figure S2a). Is the global decrease in expression burstiness in neurons simply a consequence of neurons being larger cells? To answer this question we need to first quantify the relationship between cell size and expression burstiness, as defined by the two parameters (Equations (1)-(2)).

**Figure 3:**
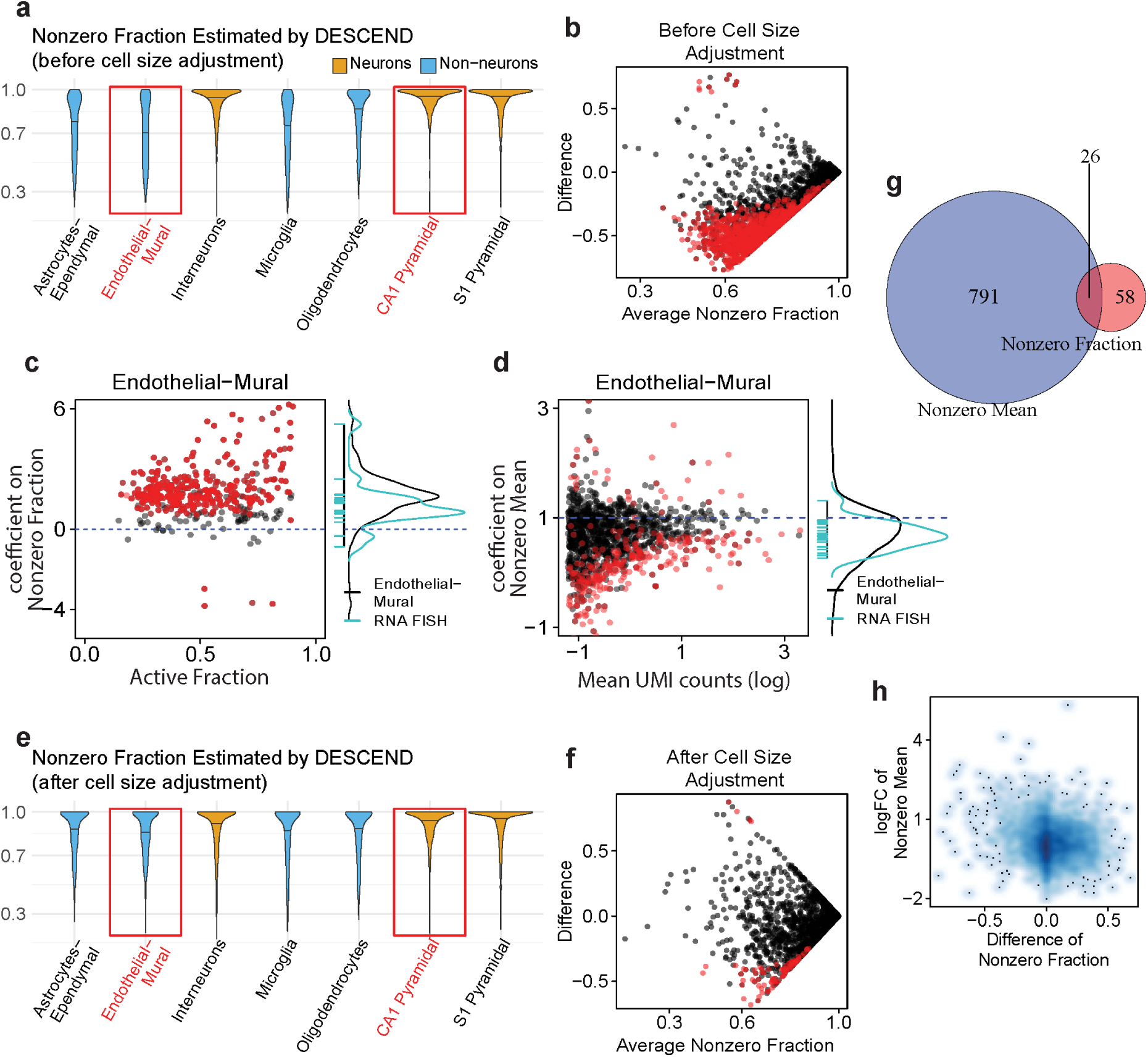
Cell size adjusted differential testing on burstiness parameters. Violin plots of the estimated nonzero fraction of the genes for each cell type (**a**)before and (**e**)after cell size adjustment. MA-plot (without log transformation) for the difference of the estimated nonzero fraction between a non-neuron cell type Endothelial-Mural and a neuron cell type CA1 Pyramidal (**b**)before and (**f**)after cell size adjustment. Red dots are the significant genes with FDR nominal level at 5%. (**c**) Left: estimated coefficients of cell size on nonzero fraction for genes whose nonzero fraction is significantly smaller than 1 and with estimated value less than 0.9 for the Endothelial-Mural cell population. Right: density of the all the dots on the left figure (black curve) aligned with the density curve of the coefficients of cell size on nonzero fraction for the RNA FISH data (blue). (**d**)Same as figure (**c**), but for coefficients on nonzero mean and all the genes. (**g**) Venn Diagram of the number of significantly differential genes on cell size adjusted nonzero mean versus on cell size adjusted nonzero fraction. (**h**)Change of nonzero fraction versus the change of nonzero mean of the genes between two cell types: Endothelial-Mural and CA1 Pyramidal.

We applied DESCEND, with cell size as a covariate, to obtain the deconvolved cell-size adjusted gene expression distribution for each gene in each of the seven cell types. Here, the cell size is defined as the total amount of RNA transcripts in each cell, which is estimated as the ratio of the library size and cell-specific efficiency. The cell efficiency, defined as the proportion of transcripts in the cell that are sequenced, is estimated from the ERCC spike-ins. The coefficients estimated by DESCEND allow us to assess, for each gene, whether its nonzero mean has super-linear, linear, or sub-linear growth with cell size, and whether its nonzero fraction increases, remains constant, or decreases with cell size. See statistical details in Methods. Taking the endothelial-mural cells as an example, we find that for almost all genes, nonzero fraction increases with cell size (Figure 3c). The mean trend across genes is that a doubling of cell size is associated with at least a doubling of the odds of observing at least one transcript. We also find that, globally, nonzero mean has a slightly sub-linear dependence on cell size, with the median scaling factor on the log scale being approximately 0.7 (Figure 3d). The sub-linear dependence of nonzero mean on cell size is consistent with previous findings in Padovan-Merhar et al. [39], which used RNA FISH to study a small set of genes and found their expression to have increased concentration in smaller cells, although the quantity measured in Padovan-Merhar et al. [39] directly reflects transcription burst size. These trends between burstiness and cell size are consistent across all seven cell types in this study (Figure S2ef).

To examine whether the estimated relationship between cell size and expression burstiness are due to unaccounted technical biases in scRNA-seq, we also quantified the relationship between cell size and expression burstiness in the RNA FISH data of Torre et al. [56]. See Supplementary Text for details. For the 23 genes in the RNA FISH data, we observe the same trends as above: the nonzero fraction increases with cell size, with a mean odds ratio of at least 2, and nonzero mean increases sub-linearly with cell size at a power of 0.7 (Figure 3cd, Figure S2c). The fact that this trend is observed under two different technologies and for eight different cell types (seven by scRNA-seq, melanoma cell line by RNA FISH) suggest that it reflects a general relationship between expression burstiness and cell size.

Figure 3e shows the nonzero fractions across genes within each cell type, estimated by applying DESCEND with cell size as a covariate. After adjusting for differences in cell size, the transcriptome-wide distribution of expression burstiness is much more similar across cell types (Figure 3e). This suggests that the increased nonzero fraction in neuron cell types can mostly be attributed to cell size differences. As an example of a cell-size-adjusted screen for genes with significant change in nonzero fraction or nonzero mean, we compared two cell types: endothelial-mural and cells pyramidal CA1 cells. Before cell size adjustment, 879 genes show significant decrease of nonzero fraction in pyramidal CA1 at FDR 5% (Figure 3b); the results change substantially after cell size adjustment, with only 84 significant genes (Figure 3f), 78 of which were in the original set of 879 genes (Figure S2b). This highlights the importance of cell size adjustment in differential testing.

Finally, we compare differential testing results on nonzero fraction, with that on nonzero mean, after adjusting for cell size. Again, we take as example endothelial-mural cells versus pyramidal CA1 cells. We detect much more genes with significant change in nonzero mean than in nonzero fraction (817 genes versus 84 genes), with only 21 genes with significant change in both (Figure 3g). Across genes, the estimated change in nonzero fraction do not exhibit any particular dependence on the estimated change in nonzero mean (Figure 3h), indicating that, after accounting for cell size, change in nonzero fraction and change in nonzero mean are independent events for most of the genes. We conducted a GO over-representation analysis of the genes that have significant increase of nonzero fraction in pyramidal CA1 cells but no significant change in nonzero mean, and found an enrichment for neuron signal transmission (Figure S3a). In contrast, a GO over-representation analysis of the 21 genes that have significant change in both nonzero fraction and nonzero mean shows enrichment of more general neuron developmental processes (Figure S3b). The differential burstiness analysis detects not only significant genes, but also gives a more detailed characterization of the nature of the change in distribution.

### DESCEND improves the accuracy of cell type identification by a better selection of highly variable genes

One major step in cell type identification is the selection of highly variable genes (HVG) before applying any dimension reduction and clustering algorithms [2, 35, 28, 59]. Filtering out genes with low variation reduces noise while keeping the genes that are more likely to be cell sub-population markers. Current pipelines select HVGs mainly by computing dispersion measurements, such as CV and Fano factor, directly on the raw or library-normalized counts, or by variance decomposition. However, as shown in Figure 2b, these methods are not as accurate as DESCEND in estimating the true biological dispersion of the gene expression levels. Furthermore, compared with CV and Fano factor, the Gini coefficient is a more robust indicator of dispersion (see Methods for derivation), and we have shown that DESCEND allows accurate estimate of this indicator. Here, we examine whether DESCEND improves the accuracy of cell type identification when combined with existing clustering algorithms.

We consider cell type identification in two datasets where somewhat reliable cell type labels are available. One consists of the 3005 cells in Zeisel et al. [62], which were classified into 7 major cell types with the help of domain knowledge. The other is obtained from ten purified cell populations derived from peripheral blood mononuclear cells (PBMC) in Zheng et al. [63], where 1000 cells were sampled from each of the ten cell types and combined to form a 10000 cell dataset. Since the PBMC data consists mostly of immune cell subtypes (CD4+ Helper T cells, CD4+/CD25+ Regulatory T Cells, CD4+/CD45RA+/CD24- Naive T cells, CD4+/CD45RO+ Memory T Cells, CD8+ Cytotoxic T cells and CD8+/CD45RA+ Naive Cytotoxic T cells) which are well-known to have very similar transcriptomes, it is a much more challenging test case. For both datasets, we treat the cell type labels given in their original papers as “gold standard”.

There are many existing and emerging cell clustering methods for scRNA-seq, but to clarify our focus of evaluating the effectiveness of the initial HVG selection step, we focus on one of the most widely used algorithms, Seurat. We compare the clustering results of Seurat (Version 2.1) with a modified version of Seurat where the initial HVG selection step is replaced by DESCEND. The original Seurat selects HVGs by computing and ranking the Fano factors of the normalized counts, yielding a list that has only around 50% overlap with the HVGs selected by DESCEND (Figure 4a) in both datasets. To compare the cell clustering accuracy, we use the adjusted Rand index, which ranges from 0, for a level of similarity expected by chance, to 1 for identical clustering [26]. The number of clusters is set to be the truth for both data sets and both pipelines. Figure 4b shows that with DESCEND, Seurat achieves consistently better results than its original version. Seurat, like most other clustering algorithms, first drastically reduces the dimension of the data using PCA, and the number of PCs chosen at this step can affect the downstream clustering result. As seen from Figure 4b, the accuracy boost obtained from DESCEND-based HVG selection is consistent across the choice of the number of principal components in the dimension reduction step.

**Figure 4:**
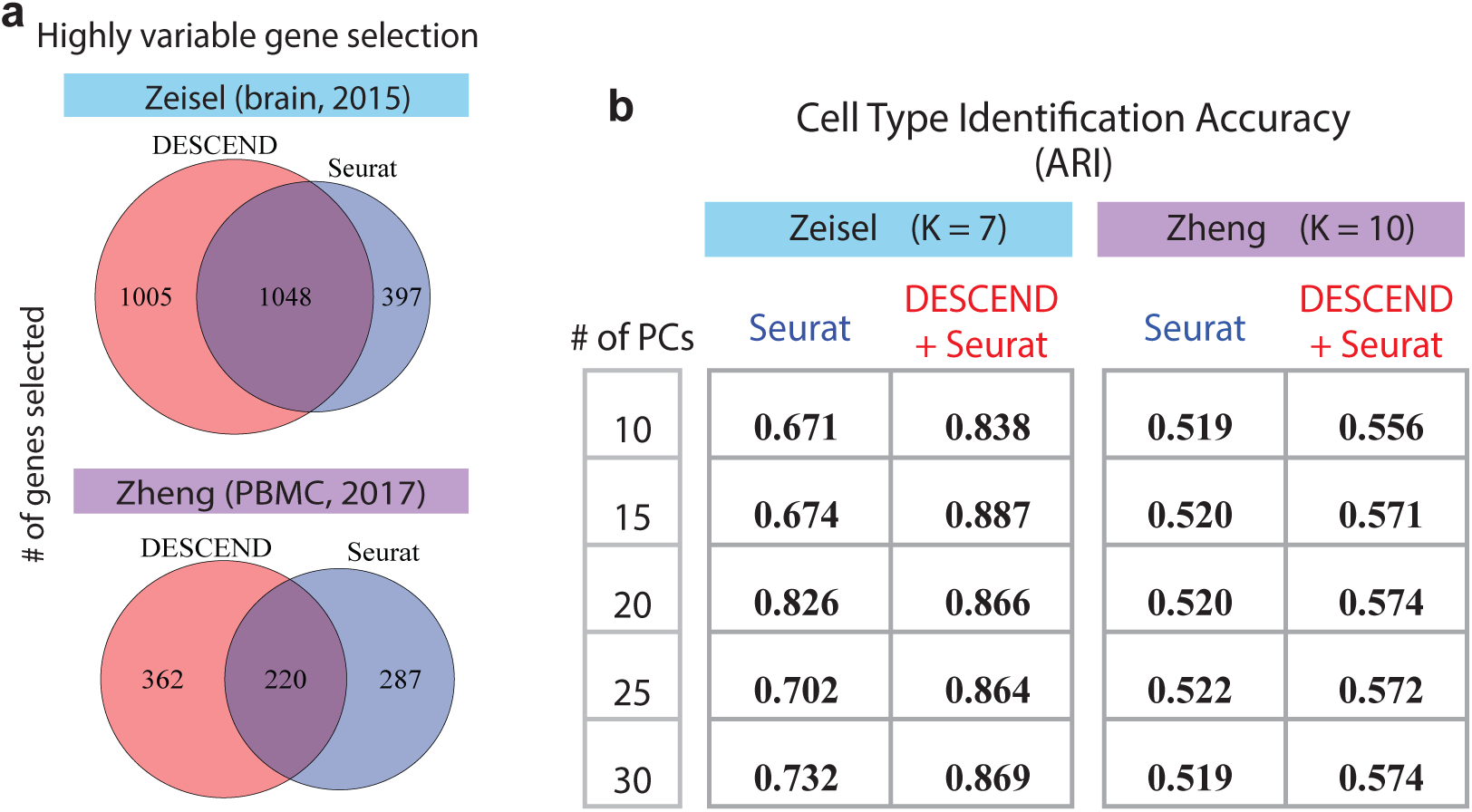
Selection of HVGs and cell type identification. (**a**) Venn Diagram of the number of selected HVGs in Seurat and using DESCEND based on Gini coefficient. (**b**)Comparison of cell type identification accuracy using ARI between the original Seurat and Seurat with HVG selection step replaced by DESCEND.

### Dispersion analysis using DESCEND estimated Gini coefficient highlights early-stage differentiation markers for mES cells

We apply DESCEND to ESC cell data from Klein et al. [28], where single cells are sampled from a differentiating mESC population before and at 2, 4, 7 days after LIF withdrawal. In this data, while pluripotency markers and differentiation-related genes have changes in mean expression over time, due to complete transcriptome remodeling, by day 7 almost all genes are significantly differentially expressed. Thus, differential analysis on mean expression is not an effective way to select differentiation markers. Instead of focusing on the mean, we test for changes in gene expression dispersion across the early time points, under the rationale that early differentiation markers would exhibit high heterogeneity across the differentiating population.

We quantify expression dispersion by the Gini coefficient computed from the DESCEND deconvolved relative gene expression distribution. This data set is complicated by the fact that the cells from days 2 and 4 are sequenced at much lower average depth than the cells from day 0 and day 7. Since low sequencing depth leads to inflation of Gini coefficient [56], Gini coefficients computed on the observed expression distributions, prior to DESCEND deconvolution, display patterns that are strictly confounded with sequencing depth (Figure S4c-e). Thus, it is especially important for this data to remove technical differences prior to drawing inferences based on the Gini coefficient, hence motivating our analysis using DESCEND.

First consider global patterns of expression dispersion. As shown in Figure 5a, the expression dispersion of most genes first increase at days 2 and 4, during differentiation, then decrease at day 7, when differentiation is mostly completed. This is expected, since during differentiation the population of cells should exhibit higher heterogeneity than either before or afterwards. However, given the much lower sequencing depth for days 2 and 4, we would also expect such increase in dispersion simply due to estimation bias. Through the DESCEND analysis, which removes technical noise, we confirmed that the increased spread in Gini at days 2/4, as compared to days 0/7, is real and not due to estimation bias. Furthermore, the correlation between the Gini estimates for day 2 and day 4, at 0.54, is as high as the correlation between the estimates for day 0 and day 7 (Figure 5b). The high correlation of the Gini coefficients between Day 0 and Day 7 is related to the fact that the cell population is mostly homogeneous at both the two days. In comparison, correlation is almost 0 between days 0/7 and days 2/4, lending confidence that DESCEND is giving biologically meaningful results.

**Figure 5:**
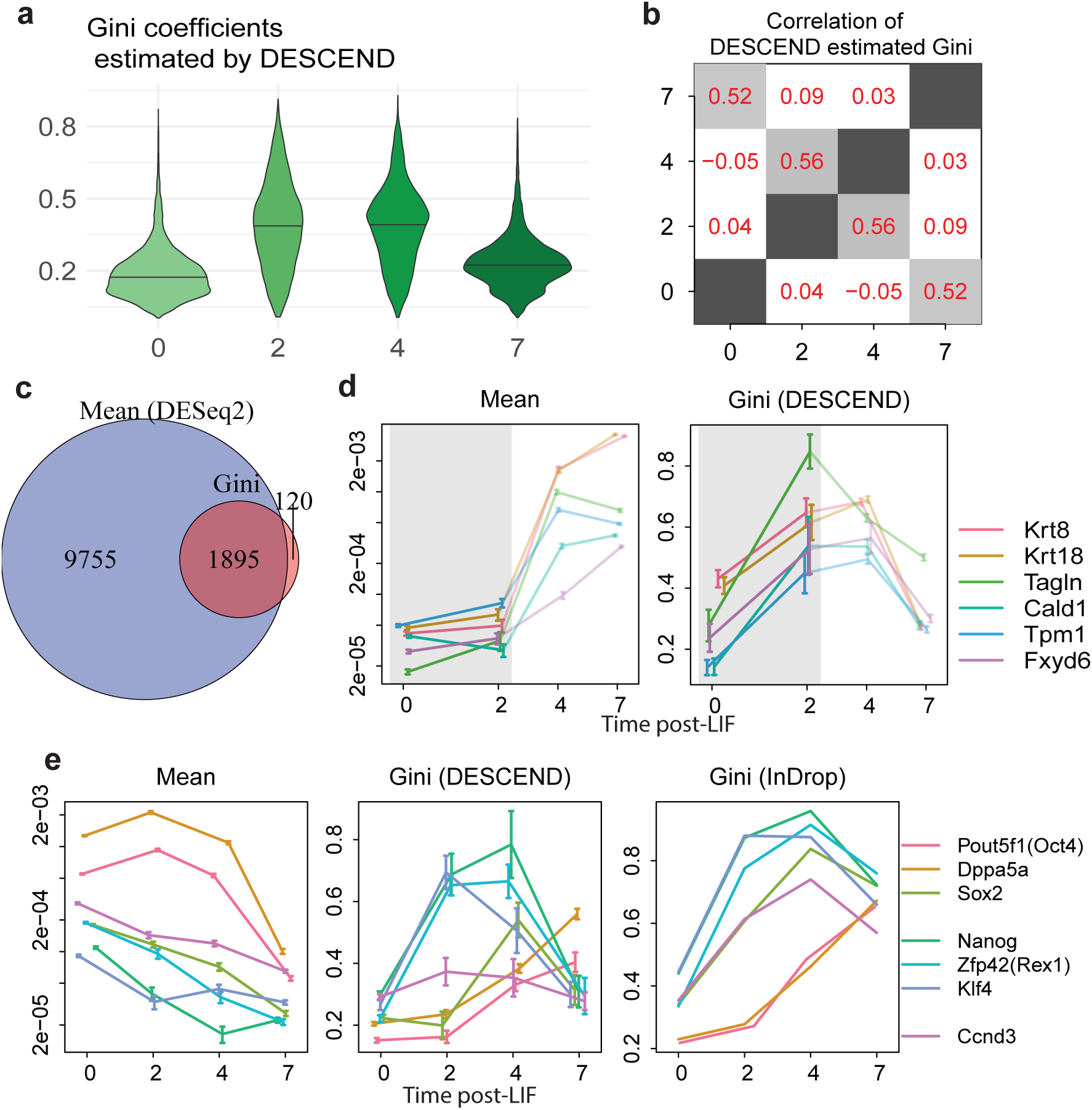
Understanding marker genes using Gini Coefficients for Klein et al. [28]. (**a**)Violin plot (with blank line indicating the 50% quantile) of the DESCEND estimated Gini coefficients for each day. (**b**)Correlation of the estimated Gini coefficient between days. (**c**)Venn diagram of differential expressed genes based on mean relative expression (tested using DESeq2) and Gini coefficient (tested using DESCEND) between Day 0 and Day 2. FDR is controlled at 5%. (**d**)Change of the mean relative expression and Gini coefficients for 6 epiblast marker genes across days. The colored error bars indicate one standard errors. (**e**)Change of the mean relative expression and Gini coefficients for 7 other genes across days. For the Gini coefficients, one is estimated using DESCEND, the other is calculated using the raw normalized counts. The colored error bars indicate one standard error.

The differentiation of mES cells upon LIF withdrawal is a poorly characterized process, with most cells committing to the epiblast lineage. Klein et al. [28] observed that, whereas at day 7, almost 100% of cells have become epiblasts, at day 2 the proportion of epiblast cells is below 10%. Thus, the cross-cell mean expression of known epiblast markers, such as Krt8, do not show significant increase at day 2. In fact, for epiblast marker genes such as Krt8, Krt18, Tagln, Cald1, Tpm1 and Fxyd6, mean expression have a substantial increase only between day 2 and day 4 (Figure 5d), when almost half of the cells and most of the transcriptome have been completely reprogrammed (Figure S5a). Thus, testing for change in mean at day 4 would not focus specifically on these marker genes. In comparison, these known marker genes belong to a much smaller set of genes that see a dramatic increase in Gini coefficient between day 0 and day 2 (Figure 5d).

DESCEND allows the computation of standard errors, and from these standard errors we assessed the significance of change in Gini coefficient between day 0 and day 2, see Methods. At an FDR threshold of 7%, 2015 genes are significant for change in Gini coefficient. In comparison, 11650 genes have significant, but very small changes, in mean between day 0 and day 2, with the significance driven mostly by the much smaller standard errors for mean estimation (Figure 5c). Of the 2015 genes with significant change in Gini coefficient, 115 genes do not have significant change in mean (Figure S4a); for these genes the GO biological process “positive regulation of epithelial cell migration” is significantly enriched (p-value 0.01, 6 out of 115 genes). Most genes in this process show significant increase in mean expression at the later stages (Day 4 or Day 7), but not at Day 2 (Figure S4b). Their increase in dispersion at day 2 suggests that cells are primed for this process early during differentiation.

DESCEND also allow a more detailed characterization of the activity of pluripotency factors during differentiation [4, 40]. As discussed in Klein et al. [28], the expression levels of some pluripotency genes drop gradually (Pou5f1, Dppa5a, Sox2) while that of other genes drop rapidly (Nanog, Zfp42, Klf4) during cell differentiation, which is revealed clearly by the trend of the DESCEND estimated Gini coefficient (Figure 5e) across time. The rapid drop-off genes react early during differentiation, and thus their Gini coefficients increase immediately at Day 2 (Figure S5). In comparison, the gradual drop-off genes react late during differentiation and thus their Gini coefficients remain unchanged at Day 2, only starting to increase at Day 4 and, for some genes, continuing to increase at Day 7. In contrast, this difference in early versus late expression drop-off is not visible by mean expression due to heteogeneity between cells with regards to their differentiation timing. As a negative control, the cell cycle marker gene Ccnd3 has no significant change in DESCEND-estimated Gini coefficient during differentiation, agreeing with the fact that the expression heterogeneity across cells for Ccnd3 should not change drastically during differentiation.

## Discussion

We have described DESCEND, a method for gene expression deconvolution for single cell RNA sequencing data. All results in this paper are based on a Poisson model for unique molecular identifier counts, which we evaluated based on ERCC spike-ins from nine published studies. The deconvolution accuracy of DESCEND was also extensively assessed through comparisons to RNA FISH performed on the same cell population, and through data splitting and simulation experiments. The effectiveness of DESCEND in removing known batch effects, as demonstrated on the data from Tung et al. [57], also testifies to the usefulness of its noise model.

DESCEND in principle can be adapted to noise models beyond the Poisson, which is necessary for analysis of scRNA-seq data without UMIs. In such data, one would need to account for amplification bias and zero-inflation beyond what could be explained by Poisson sampling [20, 58]. But such models contain many more parameters, and the estimation of these parameters from limited spike-in data is non-trivial. The estimates of expression burstiness parameters, such as nonzero fraction and nonzero mean (1, 2) are highly sensitive to the noise model and the quality of its parameter estimates. We have applied DESCEND to non-UMI data with the error model from Jia et al. [20], but the results are not satisfactory. More efforts are needed to develop robust error models for non-UMI scRNA-seq data.

Even in UMI-based scRNA-seq data, technical noise may have non-ignorable over-dispersion, and a Negative Binomial distribution may be more appropriate. DESCEND can also accept the Negative Binomial distribution with a known over-dispersion parameter *θ*. As shown in Figure 2a, *θ* can be estimated using the spike-in genes as the square of the limiting constant of CV when the expected number of input molecules are large enough. The over-dispersion parameter may also be cell or gene specific, but that would bring more complexity into the model, and a more realistic model with more parameters may not always lead to better analyses if those parameters can not be estimated reliably from the data. So far, we have settled on a simple model with at most one over-dispersion parameter for UMI-based data.

Without covariate adjustment, DESCEND currently requires a few seconds for deconvolution of the distribution of a single gene with hundreds of cells and 10 to 20 seconds when there are thousands of cells. Adding covariates and performing likelihood ratio tests can increase the computational cost roughly three or four times. Computation is easily parallelized across genes.

In general, the accuracy of DESCEND estimation increases with the number of cells and the library size. For example, for the Drop-seq data from Torre et al. [56], although the average UMI count per cell is only around 1500, the large number of cells (a few thousands) allow accurate DESCEND estimation. For the data from Klein et al. [28] and Zeisel et al. [62], there are only a few hundred cells within each condition (cell type for Zeisel et al. [62], time point for Klein et al. [28]), but the high sequencing depth and the pre-filtering allow DESCEND to make tolerably accurate estimates.

To conclude, DESCEND is a statistically rigorous and computationally efficient framework to deconvolve the true cross-cell gene expression distribution from observed scRNA-seq data. With proper use of DESCEND, one can achieve better characterization of transcriptional burstiness, gene expression stochasticity and cell population heteogeneity, and gain new biological insights from the data.

## Code availability

The R package for DESCEND is at: http://github.com/jingshuw/descend

## Online Methods

### Model

The observed count *Y_cg_* for gene *g* in cell *c* is modeled as a convolution of the true gene expression *λ_cg_* and independent technical noise:

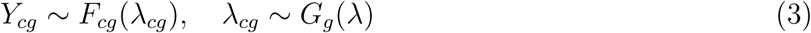

where *F_cg_*(·) quantifies technical noise and *G_g_* represents the true expression distribution of gene *g* across cells. We set *G_g_* to be a zero-inflated distribution to reflect expression burstiness. To model the effects of covariates *X_c_* on gene expression levels the gene expression *λ_cg_* is further modeled as

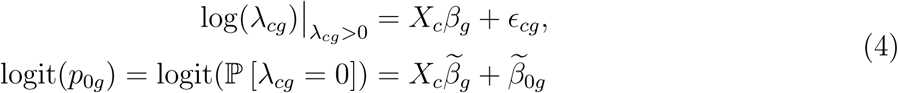

where *β_g_*, 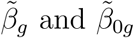 are unknown parameters, indicating that the covariates may affect both the nonzero mean and nonzero fraction.

### Estimation of the technical noise

For UMI data, we model the UMI count as

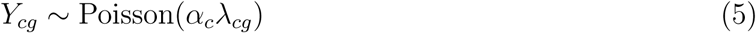

where *α_c_* is a cell specific efficiency constant. When spike-in genes are available, the efficiency is estimated as 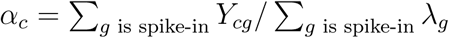 where *λ_g_* is the expected input molecule count of the spike-in genes.

When spike-in genes are not available, we recover instead the distribution of the relative expression level, defined as a gene’s expression level relative to the total quantity of RNA *N_c_* in the cell. This is equivalent to setting *X_c_* = log *N_c_* and *β_g_* = 1 in model (4). Then model (5) can be rewritten as

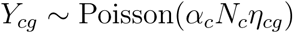

where *η_cg_* = exp(*∈_cg_*) in Equation (4) is the relative gene expression. Though neither *α_c_* nor *N_c_* is observable, their product *α_c_N_c_* can still be estimated, which is the library size 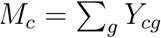 of the cell.

DESCEND also can estimate more complex technical noise models beyond (5). For example, one may assume a gene specific technical model 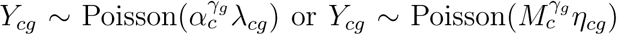 where *γ_g_* is unknown. One can simply treat *α_c_* or *M_c_* as a covariate to estimate *γ_g_*.

Another extension is to assume *Y_cg_* ∼ Negative Binomial(*α_c_λ_cg_,θ*) where *θ* is the over-dispersion parameter. *θ* can be estimated from the spike-in genes when they are available.

### Discretization of the gene expression distribution

We assume that the expression *λ_cg_* or the relative expression *η_cg_* follows a zero-inflated exponential family distribution. Following Efron (2016), the sufficient statistics of the exponential family distribution are learned from the data using natural cubic splines.

Consider the covariates adjusted model (4). Conditional on *η_cg_* > 0, we assume that *η_cg_* has density function

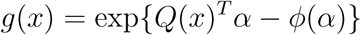

where *α* is a vector of parameters and *ϕ*(*α*) is the normalization factor. A specific form of *Q*(*x*) corresponds to a specific parametric model. For instance, when *Q*(*x*) = (log *x x*) then *g*(*x*) is the density of a Gamma distribution. When *Q*(*x*) = (log *x* (log *x*)^2^)), *g*(*x*) is the density of a logNormal distribution. In practice, we do not know which *Q*(*x*) to choose. The G-modeling idea is to set *Q*(*x*)*^T^α* as a five-degree natural cubic spline function so that the model can learn which *Q*(*x*) to use from the data.

One technique to simply the model and estimation is to discretize the distribution of *η_cg_*. Assuming that *η_cg_* can only be taken from a finite set of values ***η*** = (*η*_1_, …, *η_m_*), then we let

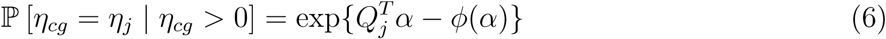

where *Q* = (*Q*_1_ … *Q_m_*)*^T^* is the five-degree natural cubic spline base matrix. In DESCEND, the default setting is *m* = 50 and we choose {*η*_1_, …, *η_m_*} as equally spaced points between 0 and the 1 — *a* percentile of {*Y_cg_*/*α_c_*, *c* = 1, 2, … *C*, *Y_cg_* ≠ 0} where *a* = 95% by default. To make sure that the choice of *η* is not affected by the covariates *X_c_*, we center *X_c_* first in our algorithm.

### Penalized maximum likelihood estimation of the model parameters

Let 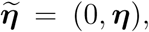 then an equivalent formulation of the discretized distribution of *η_c_* combining models (4) and (6) is

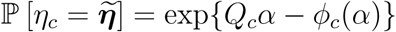

where each 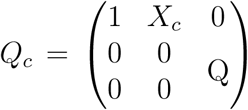 is the cell specific covariate adjusted matrix. The first two parameters in *α* can be rescaled and shifted back to 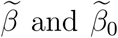 (As we recover the gene expression distribution for each gene separately, we omit the *g* subscript for notation simplicity in the following text of this section).

Let the probability of observing read count *Y_c_* as *f_c_*. Denote *p_cj_* = ℙ [*Y_c_*| *η_c_* = *η_j_*] = *F_c_*(*η_j_*) as the technical noise contribution and *g_jc_* = ℙ [*η_c_* = *η_j_*], then we have

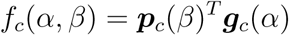

where the coefficient *β* can either be a known (when we recover the gene relative expression) or unknown coefficient. The log-likelihood of the data is ∑*_c_* log *f_c_*(*α,β*). When *β* is unknown, both *α* and *β* are parameters. When *β* is known, then only *α* is the parameter.

As suggested in Efron (2016), we maximize a penalized log-likelihood to reduce the variance of the estimation

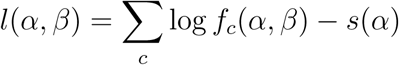

where *s*(*α*) =*c*_0_||*α*||_2_ is the penalty term. Let the Fisher information matrix of *α* be *I*(*α*). We adaptively choose the regularization constant *c_0_* such as the approximated ratio of artificial to genuine information 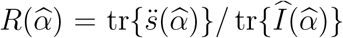 (see Efron 2016 for more details) is less than 1% to avoid over shrinkage but more than 0.05% to reduce over-fitting.

### Statistical inference

Efron (2016) showed that second-order approximation formulas provide useful inference on 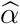 and 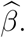 The second-order approximation uses Taylor expansion around the true value of *α* and *β*

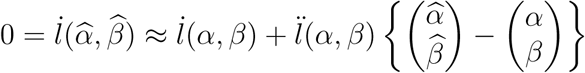

to get an estimated bias and standard deviation of our estimates using the sandwich estimator (see more details in Efron 2016). To further estimate bias and standard deviation of the estimates of the mean expression, nonzero fraction/mean, CV and Gini coefficients, we use again the second order approximations that for any function *h*(·), Taylor expansion yields 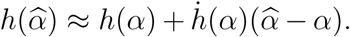. As a consequence, we get 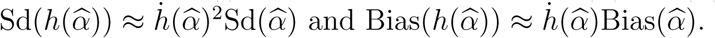.

Differential analysis of the mean expression, nonzero fraction/mean, CV and Gini Coefficients between two populations of cells are based on these approximated bias and standard deviations. Specifically, if we want to test *H*_0_ : *θ*_1_ = *θ*_2_ where *θ_i_* (*i* = 1, 2) is some parameter in population *i,* they we compute the following *Z*-score:

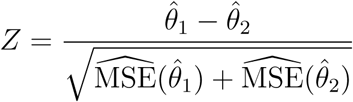

where 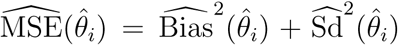 is the estimated mean squared error. We find the above Z-score construction work very well in practice.

We run permutations of the cell labels of the two populations to compute the null distribution of the Z-scores, which gives p-values [45]. Let *Y*_1_ and *Y*_2_ be two read count matrices of two cell populations. We shuffle and randomly reassign the cell samples to each population to get new shuffled read count matrices 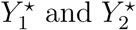 which have the same dimensions as *Y*_1_ and *Y*_2_ respectively. We compute the Z-scores *Z*\* comparing the two permuted data matrices 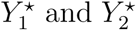 for all the genes and use the distribution of *Z*\* across genes as the null distribution of the z-scores. The p-values are then computed by calculating the quantiles of the z-scores in the null distribution.

Within on cell population, DESCEND uses likelihood ratio tests to test for specific values of nonzero fraction (for example, null hypothesis for gene *g*: ℙ [*λ_cg_* ≠ 0] = 1 testing if there is any zero inflation) and the effects of covariates on nonzero fraction and nonzero mean. For instance, we may test if cell size has a linear effect on nonzero mean (test if the coefficient of log cell size is 1 on nonzero mean) and has any effect on nonzero fraction (test if the coefficient of log cell size is 0 on nonzero fraction). For instance, to test *H*_0_ : *p*_0*g*_ = 0 vs *H*_1_ : *p*_0*g*_ > 0, we calculate the maximized unpenalized likelihood 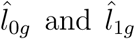 under the null and alternative hypotheses, respectively. The likelihood ratio test for gene *g* is 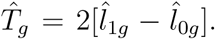 Under the null hypothesis, this test statistic approximately follows a chi-squared distribution with one degree of freedom.

### Finding highly variable genes

First, we use quantile smooth regression with a user-specific quantile *q* (default value is 50%) to fit a smooth curve (using splines) of the relationship between mean relative expression and the dispersion measurements (either CV or Gini coefficient) using R package quantreg [29]. Then, we compute the dispersion score of the each gene as the distance of the dispersion measurement from the curve and normalize it by the standard error of the dispersion measurement. Then, we select HVGs as the genes whose normalized dispersion scores are larger than a threshold *T* (default value is 10). One can reduce *T* when the experiment is not very powerful to get enough number of HVGs.

### A mathematical comparison of the CV, Gini coefficient and Fano factor

Both the CV and Gini coefficients can measure the dispersion of a distribution, and are scale invariant. Actually, their mathematical definitions are also very similar. The difference is that the Gini coefficient is more robust to outliers.

Given a collection of samples (*x*_1_,*x*_2_, …, *x_n_*), the Gini coefficient is defined as

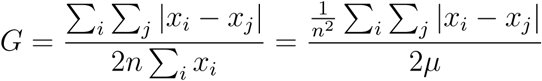

where *μ* = ∑*_i_ x_i_*/*n* is the mean of the samples. On the other hand,

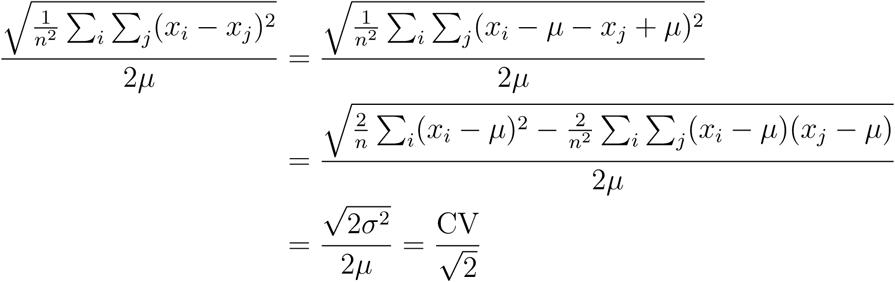

where *σ*^2^ is the sample variance. Thus, the Gini coefficient is merely a robust version of the CV as the deviations of the samples that are far away from the population center are not squared. As a consequence, we find in practice that the estimation and testing of CV and the Gini coefficients share many similarities, while Gini can be more precisely estimated in general.

In contrast, the Fano factor, defined as *σ*^2^/*μ* is not scale invariant, thus can not be estimated when the cell specific efficiency constants are unavailable. For a gene *g*, let the DESCEND recovered true gene expression be the distribution of *k_g_λ_cg_* where *k_g_* is some unknown scaling factor. Also, assume that the biological variation of gene *g* has the decomposition

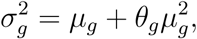

then the Fano factor of the DESCEND recovered distribution is

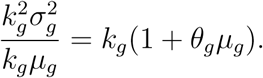

Thus, if we assume that the scaling of all the genes are approximately the same, then the ranking of the genes based on DESCEND estimated Fano factors is still reliable and favors more dispersed and highly expressed genes.

### Randomness of ERCC spike-in UMI counts in scRNA-seq experiments

The randomness of the UMI counts of ERCC spike-in genes come from two sources, one is the technical noise added through the scRNA-seq experiment, the other is the randomness of the input molecule amounts across cells.

For the technical noise, the Poisson model can be justified analytically as in Kim et al. [25]. The process in obtaining the scRNA-seq read counts contains three main steps: reserve transcription, PCR amplification and sequencing. Denote the probability of one RNA transcript being reversely transcribed as *p*_1*cg*_, then the number of copies of gene *g* in cell *c* after this step approximately follows a Binomial distribution: *X_cg_* ∼ Binomial(*λ_cg_*,*p*_1*cg*_). Also, denote the probability of one transcript being sequenced after PCR amplification as *p*_2*cg*_, the final UMI account approximately follows: *Y_cg_* ∼ Binomial(*X_cg_*,*p*_2*cg*_). Let *α_cg_* = *p*_1*cg*_*p*_2*cg*_ be the efficiency of gene *g* in cell *c*, we get *Y_cg_* ∼ Binomial(*λ_cg_*,*p*_1*cg*_*p*_2*cg*_) which is approximately Poisson(*α_cg_λ_cg_*) when the efficiency *α_cg_* is small (*Y_cg_* gets slightly less dispersed than Poisson for larger *α_cg_*). For ERCC spike-ins, by setting *α_c_* ≈ *α_cg_* we get back to our Poisson model.

For the randomness of the input molecule amounts, consider the diluted liquid of ERCC spike-ins. For the molecules of a specific spike-in gene, under the following three assumptions:

1. the molecules are distributed evenly in the liquid
2. each molecule moves independently
3. the action of taking out one drop from the liquid does not change the distribution of the molecules

we know that the number of molecules in a drop should approximately follow a Poisson distribution. If any of three assumptions is not satisfied, then the variations of the number of molecules across the drops should increase. Thus, the variation of the input molecules is at least Poisson. Thus, it is inappropriate to assume that the input number of molecules of ERCC spike-ins are constants across cells. Such randomness of the the input ERCC spike-ins is also observed empirically from the data. For example, in Zeisel et al. [62], 339 of the 3005 cells has at least one gene whose observed UMI count is even larger than the expected input molecule count, among the 38 spike-in genes that has at least 5 expected input molecule count per cell.

The randomness of the input molecule counts can be larger than Poisson if dilution is not sufficient and the assumption that each molecule moves independently fails. For example, in the ERCC dataset of Macosko et al. [35], we find out that the variances of the DESCEND deconvolved distributions roughly are 10λ where λ is the expected molecule number of a spike-in gene. The over-dispersion is high for low input values, which is reverse that of typical RNA-seq experiments where the variance changes quadratically with λ. This pattern of over-dispersion can be explained by a clumping model of the input molecular count: if 10 molecules move together on average and the molecule clusters moves randomly following Poisson, then the variance is 10λ. Thus, believe that the observed over-dispersion is more likely due to the increased randomness of input molecule instead of the inappropriate technical noise model of scRNA-seq experiment.

### Details of the parametric simulation experiment

To generate the pseudo scRNA-seq UMI counts, we utilize the observed UMI counts of 820 Oligo-dentroctyes cells from Zeisel et al. [62]. We assume that the cell size adjusted gene expression follows a zero-inflated log-Normal distribution for each gene, where the mean and variance match the corresponding parameters of the deconvolved distribution from the original observed UMI counts using DESCEND. We create “null” genes by setting the nonzero fraction as 1 for the genes whose estimated nonzero fraction using DESCEND is larger than 0.8. We keep the DESCEND estimated coefficients of cell size on nonzero mean and nonzero fraction as true parameters for this synthetic data. Only genes whose average UMI counts per cell among originally observed Oligo-dendroctyes cells is larger than 0.3 are used for generating pseudo RNA-seq counts, resulting in 5045 genes left selected. The estimated technical noise model is taken as the true technical noise model, thus the simulated UMI count data matrix has 820 cells, 5045 genes and has the same per-cell efficiency parameters and cell sizes as the original data.

We define the cell sizes as the total number of RNA copies in each cell and estimate them by the ratio between the library size per cell and the cell-specific efficiency estimated from the ERCC spike-ins. The log of cell size is added as the covariate for both the nonzero fraction and nonzero mean. We define the cell sizes of the 820 Oligodendroctyes cells as the total number of RNA copies in each cell and estimate them by the ratio between the library size per cell and the cell-specific efficiency estimated from the ERCC spike-ins. The log of cell size is added as the covariate for both the nonzero fraction and nonzero mean.

## Supplementary text and Figures

### Data sources of publicly available datasets

The ERCC UMI count matrix of Jaitin et al. [19], Macosko et al. [35], Hashimshony et al. [18] are downloaded from the NCBI GEO website (GSE54006, GSE63473, GSE78779). The raw FASTQ files of the 10x data from Svensson et al. [54] is released as ArrayExpress E-MTAB-5480 and we obtain the mapped UMI counts from the original authors. The UMI count matrices of both biological genes and ERCC spike-ins in Klein et al. [28] and Zeisel et al. [62] are downloaded from the NCBI GEO website (GSE65525, GSE60361). The count matrices of Tung et al. [57] are downloaded from the Github page: http://github.com/jdblischak/singleCellSeq. Both the ERCC data and the datasets of 10 purified cell types in Zheng et al. [63] are downloaded from the 10x genome website: http://support.10xgenomics.com/single-cell-gene-expression/datasets. To calculate the expected number of molecule amount for each ERCC spike-in gene across cells, we referred to Table 1 of Svensson et al. [54] which summarized the dilution ratios and volumes of the ERCC spike-ins in each of the publicly available datasets.

### Dispersion calculation of ERCC data from Svensson et al. [54]

For a single ERCC spike-in gene, under the Poisson noise model, as the observed count in one cell follows *Y_c_* ∼ Poisson(*α_c_λ_c_*), then conditional on *α_c_* and assume *λ_c_* ∼ Poisson(λ) where λ is the expected molecule input amount, we have Var(*Y_c_*) = *α_c_*(1 + *α_c_*)λ. as the cell efficiency *α_c_* ≈ 10^-4^ for the ERCC data from Svensson et al. [54], we have Var(*Y_c_*) ≈ *α_c_*λ and E(*Y_c_*) = *α_c_λ*.

Thus, if the Poisson noise model is correct, Var (∑_*c*_ *Y_c_*) = ∑_*c*_ *α_c_λ*. We calculate the dispersion of this spike-in gene across cells as

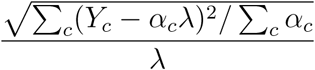

and compare it with 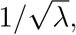 the CV of Poisson, to check if the technical noise follows Poisson or not.

The reason that we use this simple moment method instead of DESCEND to check for over-dispersion for this dataset is that under the scenario of super low cell efficiency with concentrated input molecule amount across cells (CV is mostly 0), DESCEND is biased because of the usage of a penalized likelihood and the moment method provides a more accurate estimate of dispersion for the ERCC dataset.

### Pre-processing and analysis of the RNA FISH data

The data is from Torre et al. [56] where both Drop-seq and RNA FISH are applied to the same melanoma cell line. For the Drop-seq experiment, cells with library size less than 1000 UMIs are removed (as we use GAPDH to normalize the data for recovering the relative gene expression), and 5763 cells are left with median 1473 UMIs per cell for further analysis. For the RNA FISH experiment, cells with less than 100 or more than 1000 GAPDH UMI read counts are removed, with 79099 cells left for further analysis. Of the 26 genes profiled by RNA FISH, the genes VCF and FOSL1 are removed as we find apparent inconsistency between the Drop-seq data and RNA FISH data for these two genes (The fraction of zero read counts are too high in the RNA FISH data compared with the Drop-seq data). The RNA FISH data are treated as gold standard, meaning that we ignore the measurement errors and assume that the RNA FISH counts represent the true expression level of the genes. The DESCEND recovered distribution is re-centered to have the same mean as the corresponding RNA FISH distribution as we allow for the gene-specific cell efficiency model *α_cg_* = *α_c_γ_g_*. The distribution based measurements: nonzero fraction, CV and Gini coefficients are all scaling invariant.

For its Drop-seq dataset, we computed the CV from the raw counts using conditional variance decomposition under model (5) and compare the values with the DESCEND estimated ones. For a fixed gene *g*, let *μ_Y_*(*σ_Y_*), *μ_α_*(*σ_α_*), *μ_λ_*(*σ_λ_*) be the mean (variance) of *Y_cg_*,*α_c_* and *λ_cg_* across cells respectively. Here *α_c_* is the library size of each cell. Then, we have

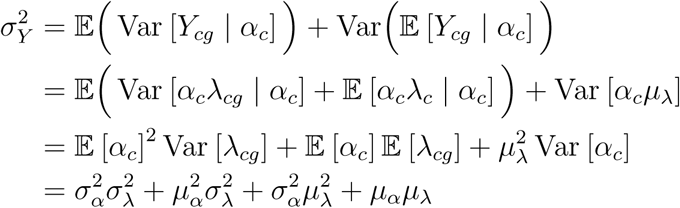

Divide by 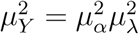 on both sides, we get

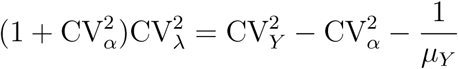

from which we can estimate 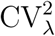 as both *Y_cg_* and *α_c_* are observable. To compute the Gini coefficient of the normalized counts, we used the R package reldist [17].

To quantify the relationship between cell size and expression burstiness in the RNA FISH data, we use the GAPDH read counts as the proxy of cell sizes, as experiments show that they are highly linearly correlated with the true cell size [39]. Thus, we have 23 genes left for further analysis (excluding VCF, FOSL1 and GAPDH). We run a logistic regression to estimated the linear relationship between the log of cell size and the odds ratio of the nonzero fraction. Also, we use R packages gamlss, gamlss.tr [46, 52] and run zero-truncated negative binomial models to estimate the linear relationship between the log of cell size and the log of the nonzero mean for the RNA FISH data.

### Analysis of the data from Tung et. al. (2017)

The data in Tung et al. (2017) contains three C1 replicates from three human induced pluripotent stem cell lines and UMI were added to all samples. In the original paper, one replicate of the first individual (NA 19098.r2) was removed from the data due to low quality and 564 cells are kept after filtering. Of the 564 cells, the number of cells in each of the eight replicates ranges from 51 to 85. For comparison between the two artificial groups, each group contains randomly selected 50 cells from one replicate of each of the three individuals. For comparison between individuals, we choose the two individuals whose all three replicates are kept and include all cells belonging to them in DESCEND.

Each replicate is a batch. As the batches are perfectly confounded with both the artificial group and the individual labels, we are unable to adjust for the confounding effects in mean expression. However, by adding the batch indicators as covariates, we can remove the batch biases within each testing group, thus can adjust for the confounding effect of batches on the dispersion parameters, such as CV and Gini coefficients.

We apply DESCEND to recover the relative gene expression and add the batches as covariates on the nonzero mean. Because of the limited number of cells in each group, we only look at the most highly expressed 187 genes whose average UMI read counts exceed 50 in order to make the estimation errors manageable. As these are highly expressed genes, most of their CV/Gini coefficients are also very small. In addition, as most of these genes are not bursty, there is no apparent difference when the batches are added as covariates on both the nonzero mean and nonzero fraction.

### Analysis of the data from Klein et al. (2015)

For the mouse embryonic stem cell data from Klein et al. (2015), the single cells are sampled from a differentiating mESC population before and at 2, 4, 7 days after LIF withdrawal. The number of cells sampled at each of the four times are 933, 303, 683 and 798 with the average library size being approximately 29500, 8500, 4700 and 26500 respectively. When comparing across all four datasets, we only keep genes whose fraction of non-zero read counts are at least 5% and whose average UMI read count are at least 0.15 in every dataset, resulting in 9059 left. For the differential analysis between Day 0 and Day 2, we ask the above criteria to be satisfied only in the two datasets involved, resulting in 13096 genes left for analysis.

In addition to using DESCEND for differential testing of the mean relative expression between Day 0 and Day 2, we also use the R package DESeq2 [34] with default settings. For the GO over-representation analysis of this dataset, we use the R package gProfileR [44]. For the tSNE plot of the differentiation at Day 4 and Day 7, we use the R package SeuratV2.1 [47] and follow the standard work flow on their online tutorial: http://satijalab.org/seurat/pbmc3k_tutorial.html.

### Analysis of the data from Zeisel et al. (2015)

The dataset contains read counts of 12234 genes in 3005 cells obtained from the mouse somatosensory cortex and hippocampus CA1 region. The 3005 cells have been further clustered into 7 major cell types: Astrocytes-Ependymal, Endothelial-Mural, Interneurons, Microglia, Oligodendrocytes, CA1 pyramidal and S1 pyramidal, and the number cells in each cell type are 224, 235, 290, 98, 820, 939 and 399 respectively. We recover the cell size adjusted gene expression distribution for each cell type separately. The cell sizes are estimated as the ratio between the library size and the cell efficiency estimated from the ERCC spike-ins. To avoid estimation bias, both the cell efficiency and cell size are treated as covariates on both nonzero fraction and nonzero mean.

For each cell type, we only recover the expression distribution of the genes whose fraction of non-zero read counts are at least 5% and whose average UMI read count are at least 0.3 for estimating the burstiness parameters with acceptable accuracy. The number of genes kept are then 3855, 3496, 7984, 3299, 4951, 7866 and 7683 respectively for each cell type. When we compare across cell types, we only consider the intersection of the genes from the involved cell types. For example, there are 2105 genes which are kept in all cell types.

For the GO over-representation analysis of this dataset, we use the R package clusterProfiler [61] which allows user-defined list of the background genes. We define the list of background genes to be all genes who passes our filtering criteria to avoid possible biases introduced during the filtering process.

### Cell type identification

For cell clustering of the datasets from Zeisel et al. [62] and Zheng et al. [63], we follow the steps in the online tutorial of Seurat V2.1. We try different values of the resolution parameter to get the desired number of clusters (*K* = 7 for Zeisel et al. [62] and *K* =10 for Zheng et al. [63]) in each scenario and compare the clustering accuracy of Seurat with and without DESCEND at the given number of clusters. To compute the adjusted Rand index, we use the R package mclust [13].

## Supplementary figures

**Figure S1:**
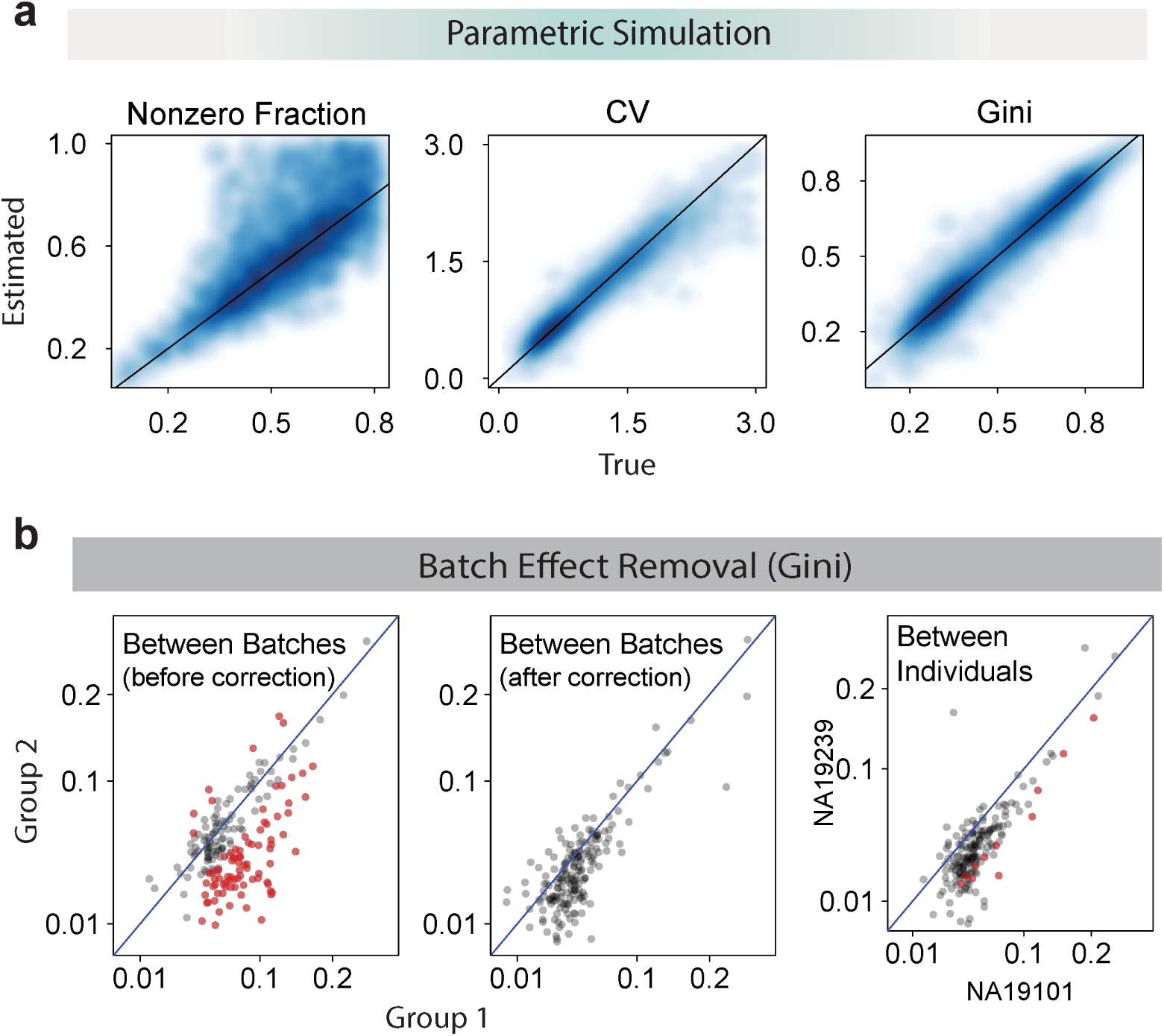
More figures for validation of DESCEND. (**a**)Comparison between the estimated nonzero fraction, CV and Gini coefficients and the true values in the parametric simulation experiment. (**b**)Batch effect removal: the DESCEND estimated Gini coefficients are compared between the two artificial groups before (left) and after (middle) adding batches as covariates. Each dot is a gene. The estimated Gini after adding batches as covariates are also compared between two individuals (right). The red dots are significantly differentially expressed genes (of CV) when FDR in controlled at level 5%.

**Figure S2:**
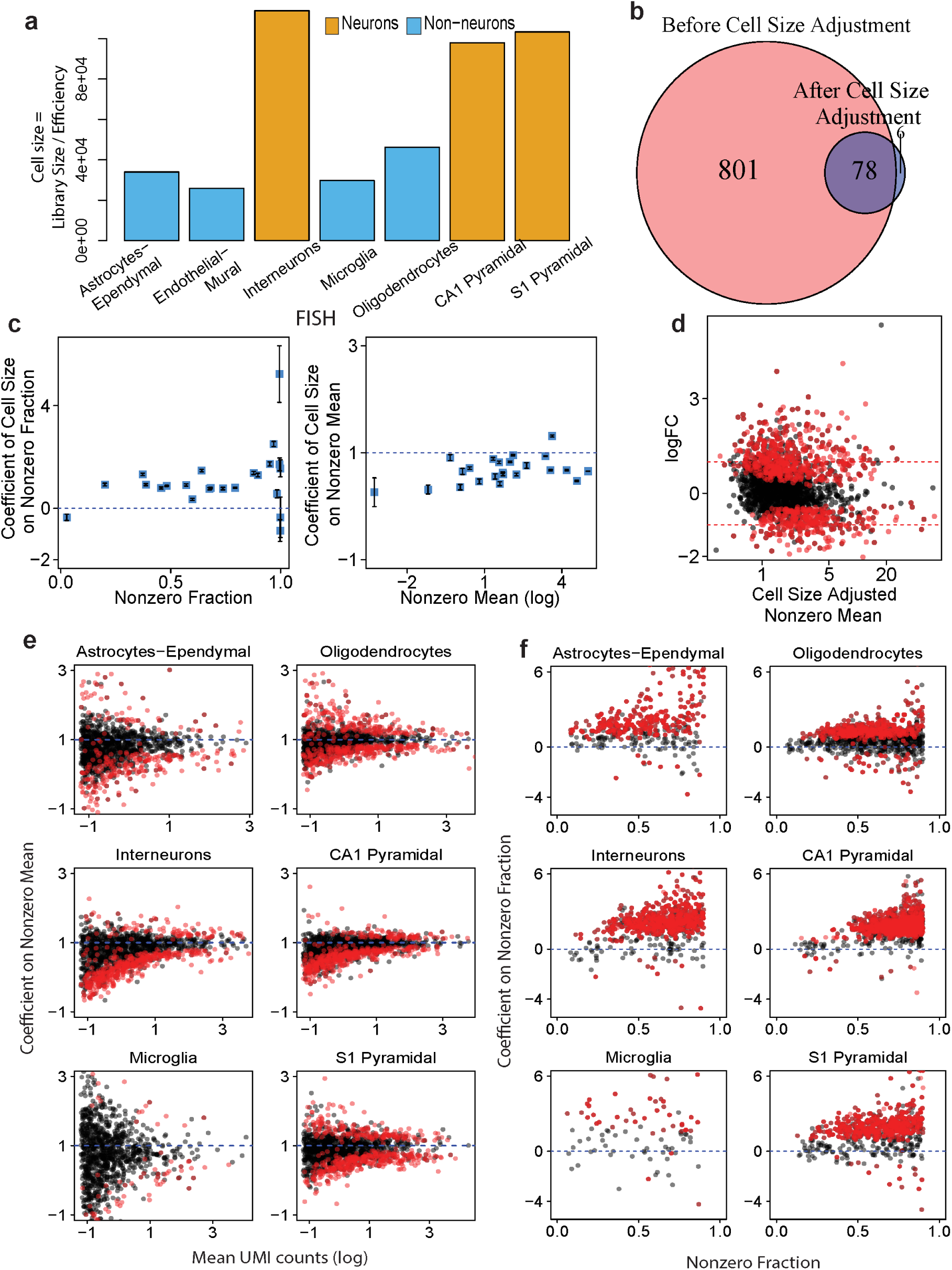
Supplementary figures for the case study of Zeisel et al. [62]. **(a)**Bar plot of the mean cell size (calculated as the ratio of library size and cell efficiency) of each cell type. Neuron cells are much larger than non-neuron cells. **(b)**Venn Diagram of the number of significantly differentially expressed genes based on nonzero fraction before and after cell size adjustment. **(c)**The coefficients of cell size on nonzero fraction and nonzero mean for the RNA FISH data. The black bars are one standard error bars. **(d)** MA plot of the estimated nonzero mean after cell size adjustment. **(e)**A consistent sub-linear effect of cell size on nonzero mean across all cell types. **(f)**A consistent positive effect of cell size on nonzero fraction across all cell types.

**Figure S3:**
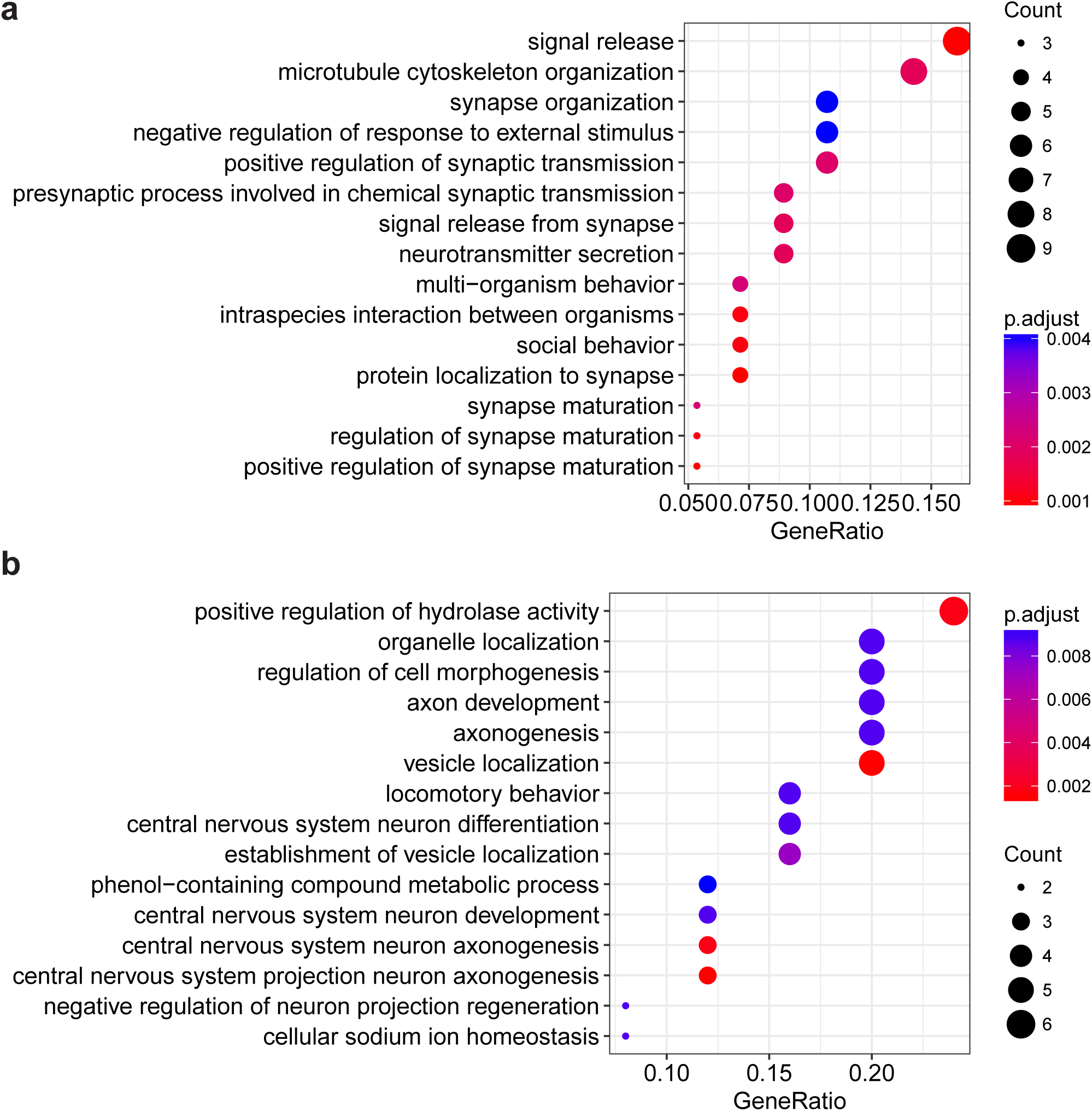
GO over-representation analysis for **(a)**genes whose nonzero fraction is significant differentially expressed but not the nonzero mean and **(b)**genes whose both nonzero fraction and nonzero mean are differentially expressed genes based on nonzero fraction between the non-neuron cell type Endothelial-Mural and the neuron cell type CA1 Pyramidal. The plot shows the 15 categories with the smallest p-values using Fisher’s exact test.

**Figure S4:**
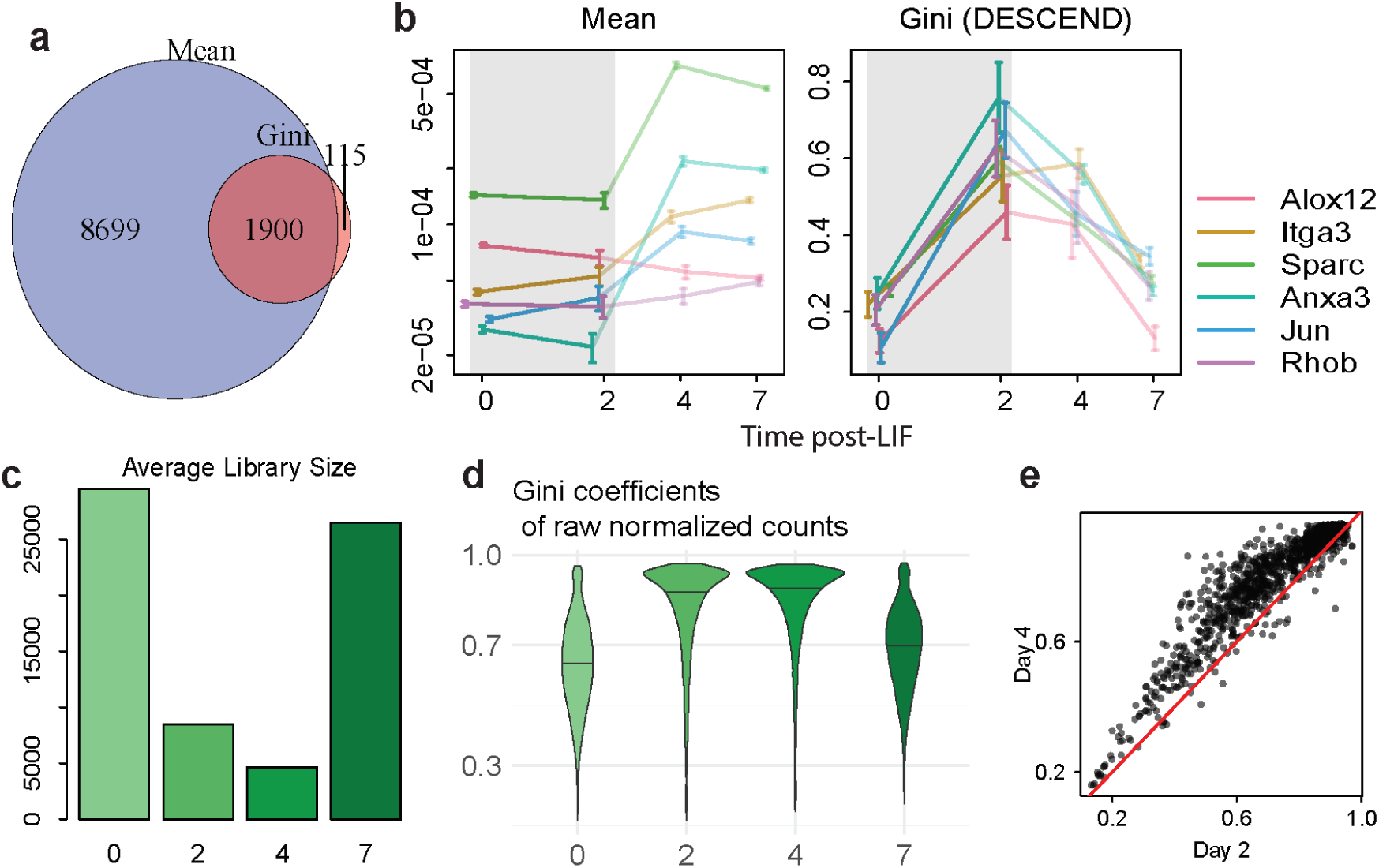
Supplementary figures for the case study of Klein et al. [28]. (**a**)Venn diagram of the number of differential expressed genes based on mean relative expression and Gini coefficient, both tested using DESCEND. (**b**)Change of the mean relative expression and Gini coefficients for 6 marker genes that contribute to the enrichment of GO term “positive regulation of epithelial cell migration”. The colored bars are one standard error bars. (**c**)Average library sizes for the cells at each day. (**d**)Violin plot of Gini coefficients directly calculated using the normalized UMI counts. (**e**)Comparison of gini coefficients directly calculated using the normalized UMI counts between day 2 and day 4.

**Figure S5:**
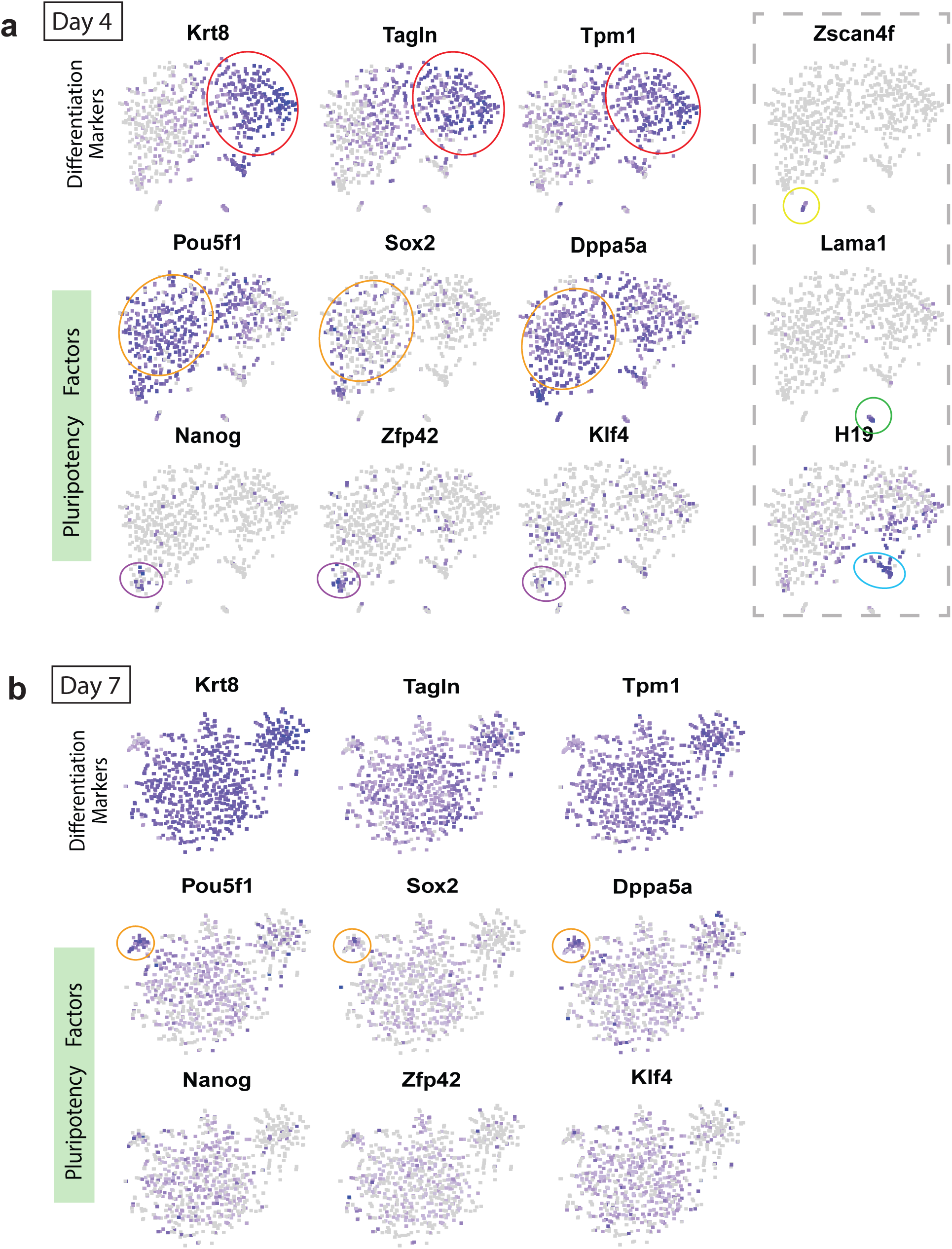
Feature plots of the tSNE plots for the cells at day 4 and day 7. (**a**)The colored circles represents the differentiated cells (red), not fully differentiated cells (orange) and undifferentiated cells (purple). There are also three other cell clusters at Day 4 (yellow, green and blue circles). (**b**) The orange circled cells are those that have not fully differentiated at Day 7.

## References

[1] R. Bacher and C. Kendziorski. Design and computational analysis of single-cell RNA-sequencing experiments. Genome biology, 17(1):63, 2016.

[2] P. Brennecke, S. Anders, J. K. Kim, A. A. Kołodziejczyk, X. Zhang, V. Proserpio, B. Baying, V. Benes, S. A. Teichmann, J. C. Marioni, and M. G. Heisler. Accounting for technical noise in single-cell RNA-seq experiments. Nature methods, 10(11):1093–1095, 2013.

[3] F. Buettner, K. N. Natarajan, F. P. Casale, V. Proserpio, A. Scialdone, F. J. Theis, S. A. Teichmann, J. C. Marioni, and O. Stegle. Computational analysis of cell-to-cell heterogeneity in single-cell RNA-sequencing data reveals hidden subpopulations of cells. Nature biotechnology, 33(2):155–160, 2015.

[4] L.-F. Chu, N. Leng, J. Zhang, Z. Hou, D. Mamott, D. T. Vereide, J. Choi, C. Kendziorski, R. Stewart, and J. A. Thomson. Single-cell RNA-seq reveals novel regulators of human embryonic stem cell differentiation to definitive endoderm. Genome biology, 17(1):173, 2016.

[5] J. R. Chubb, T. Trcek, S. M. Shenoy, and R. H. Singer. Transcriptional pulsing of a developmental gene. Current biology, 16(10):1018–1025, 2006.

[6] R. D. Dar, B. S. Razooky, A. Singh, T. V. Trimeloni, J. M. McCollum, C. D. Cox, M. L. Simpson, and L. S. Weinberger. Transcriptional burst frequency and burst size are equally modulated across the human genome. Proceedings of the National Academy of Sciences, 109(43):17454–17459, 2012.

[7] M. Delmans and M. Hemberg. Discrete distributional differential expression (D3E)-a tool for gene expression analysis of single-cell RNA-seq data. BMC bioinformatics, 17(1):110, 2016.

[8] Q. Deng, D. Ramsköld, B. Reinius, and R. Sandberg. Single-cell RNA-seq reveals dynamic, random monoallelic gene expression in mammalian cells. Science, 343(6167):193–196, 2014.

[9] J. Eberwine, J.-Y. Sul, T. Bartfai, and J. Kim. The promise of single-cell sequencing. Nature methods, 11(1):25–27, 2014.

[10] B. Efron. Empirical bayes deconvolution estimates. Biometrika, 103(1):1–20, 2016.

[11] A. Eldar and M. B. Elowitz. Functional roles for noise in genetic circuits. Nature, 467(7312): 167, 2010.

[12] G. Finak, A. McDavid, M. Yajima, J. Deng, V. Gersuk, A. K. Shalek, C. K. Slichter, H. W. Miller, M. J. McElrath, M. Prlic, P. S. Linsley, and R. Gottardo. Mast: a flexible statistical framework for assessing transcriptional changes and characterizing heterogeneity in single-cell RNA sequencing data. Genome biology, 16(1):278, 2015.

[13] C. Fraley, A. E. Raftery, T. B. Murphy, and L. Scrucca. mclust Version 4 for R: Normal Mixture Modeling for Model-Based Clustering, Classification, and Density Estimation, 2012.

[14] K. Geiler-Samerotte, C. Bauer, S. Li, N. Ziv, D. Gresham, and M. Siegal. The details in the distributions: why and how to study phenotypic variability. Current opinion in biotechnology, 24(4):752–759, 2013.

[15] D. Grün, L. Kester, and A. Van Oudenaarden. Validation of noise models for single-cell transcriptomics. Nature methods, 11(6):637–640, 2014.

[16] J. Gu, Q. Du, X. Wang, P. Yu, and W. Lin. Sphinx: modeling transcriptional heterogeneity in single-cell RNA-seq. bioRxiv, page 027870, 2015.

[17] M. S. Handcock. Relative Distribution Methods. Los Angeles, CA, 2016. URL https://CRAN.R-project.org/package=reldist. Version 1.6-6. Project home page at url-http://www.stat.ucla.edu/handcock/RelDist.

[18] T. Hashimshony, N. Senderovich, G. Avital, A. Klochendler, Y. de Leeuw, L. Anavy, D. Gennert, S. Li, K. J. Livak, O. Rozenblatt-Rosen, Y. Dor, A. Regev, and I. Yanai. CEL-Seq2: sensitive highly-multiplexed single-cell rna-seq. Genome biology, 17(1):77, 2016.

[19] D. A. Jaitin, E. Kenigsberg, H. Keren-Shaul, N. Elefant, F. Paul, I. Zaretsky, A. Mildner, N. Cohen, S. Jung, A. Tanay, and I. Amit. Massively parallel single-cell RNA-seq for markerfree decomposition of tissues into cell types. Science, 343(6172):776–779, 2014.

[20] C. Jia, D. Kelly, J. Kim, M. Li, and N. Zhang. Accounting for technical noise in single-cell rna sequencing analysis. bioRxiv, page 116939, 2017.

[21] Y. Jiang, N. R. Zhang, and M. Li. SCALE: modeling allele-specific gene expression by singlecell RNA sequencing. Genome biology, 18(1):74, 2017.

[22] M. Kaern, T. C. Elston, W. J. Blake, and J. J. Collins. Stochasticity in gene expression: from theories to phenotypes. Nature reviews. Genetics, 6(6):451, 2005.

[23] P. V. Kharchenko, L. Silberstein, and D. T. Scadden. Bayesian approach to single-cell differential expression analysis. Nature methods, 11(7):740–742, 2014.

[24] J. K. Kim and J. C. Marioni. Inferring the kinetics of stochastic gene expression from single-cell RNA-sequencing data. Genome biology, 14(1):R7, 2013.

[25] J. K. Kim, A. A. Kolodziejczyk, T. Ilicic, S. A. Teichmann, and J. C. Marioni. Characterizing noise structure in single-cell RNA-seq distinguishes genuine from technical stochastic allelic expression. Nature communications, 6:8687, 2015.

[26] V. Y. Kiselev, K. Kirschner, M. T. Schaub, T. Andrews, A. Yiu, T. Chandra, K. N. Natarajan, W. Reik, M. Barahona, A. R. Green, and M. Hamberg. SC3: consensus clustering of single-cell RNA-seq data. Nature methods, 2017.

[27] T. Kivioja, A. Vähärautio, K. Karlsson, M. Bonke, M. Enge, S. Linnarsson, and J. Taipale. Counting absolute numbers of molecules using unique molecular identifiers. Nature methods, 9(1):72–74, 2012.

[28] A. M. Klein, L. Mazutis, I. Akartuna, N. Tallapragada, A. Veres, V. Li, L. Peshkin, D. A. Weitz, and M. W. Kirschner. Droplet barcoding for single-cell transcriptomics applied to embryonic stem cells. Cell, 161(5):1187–1201, 2015.

[29] R. Koenker. quantreg: Quantile Regression, 2017. URL https://CRAN.R-project.org/package=quantreg. R package version 5.34.

[30] A. A. Kolodziejczyk, J. K. Kim, V. Svensson, J. C. Marioni, and S. A. Teichmann. The technology and biology of single-cell RNA sequencing. Molecular cell, 58(4):610–620, 2015.

[31] K. D. Korthauer, L.-F. Chu, M. A. Newton, Y. Li, J. Thomson, R. Stewart, and C. Kendziorski. A statistical approach for identifying differential distributions in single-cell RNA-seq experiments. Genome biology, 17(1):222, 2016.

[32] S. F. Levy, N. Ziv, and M. L. Siegal. Bet hedging in yeast by heterogeneous, age-correlated expression of a stress protectant. PLoS biology, 10(5):e1001325, 2012.

[33] A. Loewer and G. Lahav. We are all individuals: causes and consequences of non-genetic heterogeneity in mammalian cells. Current opinion in genetics & development, 21(6):753–758, 2011.

[34] M. I. Love, W. Huber, and S. Anders. Moderated estimation of fold change and dispersion for RNA-seq data with DESeq2. Genome biology, 15(12):550, 2014.

[35] E. Z. Macosko, A. Basu, R. Satija, J. Nemesh, K. Shekhar, M. Goldman, I. Tirosh, A. R. Bialas, N. Kamitaki, E. M. Martersteck, et al. Highly parallel genome-wide expression profiling of individual cells using nanoliter droplets. Cell, 161(5):1202–1214, 2015.

[36] C. P. Martinez-Jimenez, N. Eling, H.-C. Chen, C. A. Vallejos, A. A. Kolodziejczyk, F. Connor, L. Stojic, T. F. Rayner, M. J. Stubbington, S. A. Teichmann, et al. Aging increases cell-to-cell transcriptional variability upon immune stimulation. Science, 355(6332):1433–1436, 2017.

[37] A. McDavid, G. Finak, P. K. Chattopadyay, M. Dominguez, L. Lamoreaux, S. S. Ma, M. Roederer, and R. Gottardo. Data exploration, quality control and testing in single-cell qPCR-based gene expression experiments. Bioinformatics, 29(4):461–467, 2012.

[38] M. Narayanan, A. J. Martins, and J. S. Tsang. Robust inference of cell-to-cell expression variations from single-and k-cell profiling. PLoS computational biology, 12(7):e1005016, 2016.

[39] O. Padovan-Merhar, G. P. Nair, A. G. Biaesch, A. Mayer, S. Scarfone, S. W. Foley, A. R. Wu, L. S. Churchman, A. Singh, and A. Raj. Single mammalian cells compensate for differences in cellular volume and DNA copy number through independent global transcriptional mechanisms. Molecular cell, 58(2):339–352, 2015.

[40] D. Papatsenko, H. Xu, A. Ma’ayan, and I. Lemischka. Quantitative approaches to model pluripotency and differentiation in stem cells. In Stem Cells Handbook, pages 59–74. Springer, 2013.

[41] S. Prabhakaran, E. Azizi, A. Carr, and D. Pe’er. Dirichlet process mixture model for correcting technical variation in single-cell gene expression data. In International Conference on Machine Learning, pages 1070–1079, 2016.

[42] A. Raj and A. van Oudenaarden. Single-molecule approaches to stochastic gene expression. Annual review of biophysics, 38:255–270, 2009.

[43] A. Raj, C. S. Peskin, D. Tranchina, D. Y. Vargas, and S. Tyagi. Stochastic mRNA synthesis in mammalian cells. PLoS biology, 4(10):e309, 2006.

[44] J. Reimand, R. Kolde, and T. Arak. gProfileR: Interface to the ‘g:Profiler’ Toolkit, 2016. URL https://CRAN.R-project.org/package=gProfileR. R package version 0.6.1.

[45] A. Reiner, D. Yekutieli, and Y. Benjamini. Identifying differentially expressed genes using false discovery rate controlling procedures. Bioinformatics, 19(3):368–375, 2003.

[46] R. A. Rigby and D. M. Stasinopoulos. Generalized additive models for location, scale and shape,(with discussion). Applied Statistics, 54:507–554, 2005.

[47] R. Satija, A. Butler, and P. Hoffman. Seurat: Tools for Single Cell Genomics, 2017. URL https://CRAN.R-project.org/package=Seurat. R package version 2.1.0.

[48] S. M. Shaffer, M. C. Dunagin, S. R. Torborg, E. A. Torre, B. Emert, C. Krepler, M. Beqiri, K. Sproesser, P. A. Brafford, M. Xiao, et al. Rare cell variability and drug-induced reprogramming as a mode of cancer drug resistance. Nature, 546(7658):431–435, 2017.

[49] A. K. Shalek, R. Satija, X. Adiconis, R. S. Gertner, J. T. Gaublomme, R. Raychowdhury, S. Schwartz, N. Yosef, C. Malboeuf, D. Lu, et al. Single-cell transcriptomics reveals bimodality in expression and splicing in immune cells. Nature, 498(7453):236, 2013.

[50] A. K. Shalek, R. Satija, J. Shuga, J. J. Trombetta, D. Gennert, D. Lu, P. Chen, R. S. Gertner, J. T. Gaublomme, N. Yosef, et al. Single cell RNA Seq reveals dynamic paracrine control of cellular variation. Nature, 510(7505):363, 2014.

[51] S. L. Spencer, S. Gaudet, J. G. Albeck, J. M. Burke, and P. K. Sorger. Non-genetic origins of cell-to-cell variability in trail-induced apoptosis. Nature, 459(7245):428, 2009.

[52] M. Stasinopoulos and B. Rigby. gamlss.tr: Generating and Fitting Truncated ‘gamlss.family’ Distributions, 2016. URL https://CRAN.R-project.org/package=gamlss.tr. R package version 5.0-0.

[53] O. Stegle, S. A. Teichmann, and J. C. Marioni. Computational and analytical challenges in single-cell transcriptomics. Nature reviews. Genetics, 16(3):133, 2015.

[54] V. Svensson, K. N. Natarajan, L.-H. Ly, R. J. Miragaia, C. Labalette, I.C. Macaulay, A. Cvejic, and S. A. Teichmann. Power analysis of single-cell RNA-sequencing experiments. Nature methods, 2017.

[55] S. Tay, J. J. Hughey, T. K. Lee, T. Lipniacki, S. R. Quake, and M. W. Covert. Single-cell NF-κB dynamics reveal digital activation and analog information processing in cells. Nature, 466(7303):267, 2010.

[56] E. A. Torre, H. Dueck, S. Shaffer, J. Gospocic, R. Gupte, R. Bonasio, J. Kim, J. Murray, and A. Raj. A comparison between single cell RNA sequencing and single molecule RNA FISH for rare cell analysis. bioRxiv, page 138289, 2017.

[57] P.-Y. Tung, J. D. Blischak, C. J. Hsiao, D. A. Knowles, J. E. Burnett, J. K. Pritchard, and Y. Gilad. Batch effects and the effective design of single-cell gene expression studies. Scientific reports, 7:39921, 2017.

[58] C. A. Vallejos, J. C. Marioni, and S. Richardson. BASiCS: Bayesian analysis of single-cell sequencing data. PLoS computational biology, 11(6):e1004333, 2015.

[59] C. A. Vallejos, D. Risso, A. Scialdone, S. Dudoit, and J. C. Marioni. Normalizing single-cell RNA sequencing data: challenges and opportunities. Nature methods, 2017.

[60] T. N. Vu, Q. F. Wills, K. R. Kalari, N. Niu, L. Wang, M. Rantalainen, and Y. Pawitan. Beta-Poisson model for single-cell RNA-seq data analyses. Bioinformatics, 32(14):2128–2135, 2016.

[61] G. Yu, L.-G. Wang, Y. Han, and Q.-Y. He. clusterprofiler: an r package for comparing biological themes among gene clusters. OMICS: A Journal of Integrative Biology, 16(5):284–287, 2012. doi: 10.1089/omi.2011.0118.

[62] A. Zeisel, A. B. Muñoz-Manchado, S. Codeluppi, P. Lönnerberg, G. La Manno, A. Juréus, S. Marques, H. Munguba, L. He, C. Betsholtz, et al. Cell types in the mouse cortex and hippocampus revealed by single-cell RNA-seq. Science, 347(6226):1138–1142, 2015.

[63] G. X. Zheng, J. M. Terry, P. Belgrader, P. Ryvkin, Z. W. Bent, R. Wilson, S. B. Ziraldo, T. D. Wheeler, G. P. McDermott, J. Zhu, et al. Massively parallel digital transcriptional profiling of single cells. Nature communications, 8:14049, 2017.

